# Inactivation of Vascular Stem Cells Suppresses Intimal Hyperplasia in Vein Grafts

**DOI:** 10.1101/2022.05.17.492289

**Authors:** Weiwei Wu, Huimei Zang, Lei Qi, Mitzi Nagarkatti, Prakash Nagarkatti, Isidro Sánez-Garcia, Mary C.M Weiser-Evans, Taixing Cui

**Affiliations:** Department of Cell Biology and Anatomy, University of South Carolina, Columbia, SC 29208, USA; Department of Pathology, Microbiology and Immunology, School of Medicine, University of South Carolina, Columbia, SC 29208, USA; Experitmental Therapeutics and Translational Oncology Program, Instituto de Biología Molecular y Celular del Cáncer, Consejo Superior de InvestigacionesCientificas/Universidad de Salamanca, 37007 Salamanca, Spain; Institute for Biomedical Research of Salamanca (IBSAL), Salamanca, Spain; Department of Medicine, Division of Renal Diseases and Hypertension, University of Colorado Anschutz Medical Campus, Aurora, CO 80045, USA

## Abstract

Chronic vein graft (VG) failure (VGF) is associated with VG intimal hyperplasia characterized by abnormal proliferation and accumulation of vascular smooth muscle cells (SMCs), which are derived predominantly from pre-existing vascular SMCs via a process of vascular SMC dedifferentiation.^1^ Therapeutic approaches focused on vascular SMCs, however, have not changed chronic VGF rates,^2^ indicating additional and yet unrecognized causes of chronic VGF. Herein, we uncover a novel diagram in which after transplantation, recipient vascular stem cells (VSCs) expressing Sca1 are activated and accumulated in the adventitia of VGs, and they do not differentiate into SMCs but promote intimal hyperplasia via paracrine enforcement of medial SMC dedifferentiation into synthetic SMCs in VGs, suggesting that VSCs may be a novel target for the treatment of chronic VGF.

## Results

In normal murine jugular veins, we found that most of the VSC marker Sca1 positive (Sca1^+^) or CD34^+^ cells reside mainly in the intima and the adventitia (Figures 1a, S1, and Table S1, S4). However, the Gli1^+^cells are located predominantly in the adventitia (Figure S1 and Table S1). The heterogeneity of Sca1^+^ cells is identified by existence of substantial Sca1 and vWF double positive (Sca1^+^vWF^+^) or Scal^+^CD34^+^cells in the intima as well as Sca1^+^CD34^+^ and Sca1^+^Gli1^+^ cells in the adventitia (Figure S2 and Table S2).

**Figure 1.**
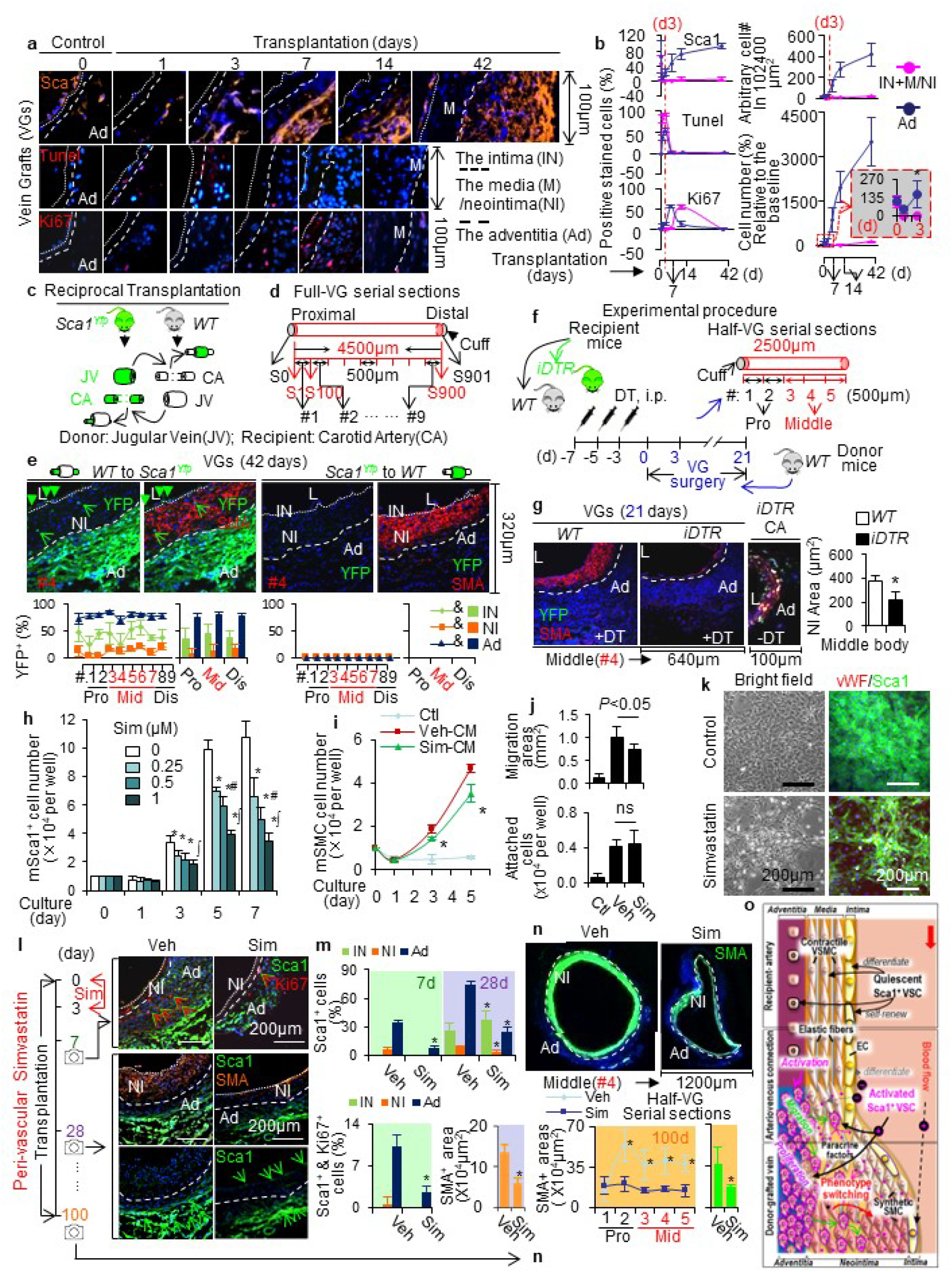
**a,** VSC dynamics in jugular vein isografts. Representative immunofluorescence staining for Sca1, Tnnel, Ki67. Small-dotted lines separate the intima (IN) and the media (M) & the neointima (NI); big dotted lines separate M&NI and the adventitia (Ad). All mice used for vein graft (VG) transplantation were male 3-month-old wild type mice in a C57BL/6J genetic background; and vessels were harvested at different time (0, 1, 3, 7, 14, 42 days) after transplantation. **b,** Quantified data of positively stained cells in 2 different layers (IN+M/NI and Ad). Four whole vessel cross-sections were randomly chosen at each time point per VG (n=8) and subject to the immunofluorescence staining analysis. All images which could be used for analysis were counted. #, numbers. **c,** A scheme of reciprocal VG transplantation. Jugular veins (JVs) were transplanted to the carotid arteries (CAs) between in wild type (*WT*) mice and *Sca1^Yfp^* mice which Sca1^+^ cells were genetically labeled with YFP. **d,** A scheme of consecutive tissue sectioning of VGs. S, VG cross-section. #1 is one of the first 100 Ss from the proximal end of VGs; #9 is one of 100 Ss of the distal end of VGs. **e,** Characterization of reciprocal VG remodeling. The representative images of co-staining of YFP and SMA in the cross-sections of #3-#7 segments at the middle part (Mid). YFP is green; SMA is red. L, the lumen; IN, the intima; NI, the neointima; Ad, the adventitia. Dotted lines separate the IN and NI as well as the NI and Ad layers. The percentages of SMA, YFP and SMA and YFP double positive cells in each major layer of VGs with two reciprocal VG-remodeling (Total mice number of each type of VG is 11 and 12, each mouse has 9 segments, each segment has 3 cross-sections). Data are means ± SEM. **f,** Scheme of experimental procedures and Half-VG serial sections. *iDTR, Sca1-Cre::R26-eYfp^fl/-^::R26-DTR^fl/-^*; DT, diphtheria toxin. **g,** The effect of genetic depletion of Sca1^+^ cells on VG remodeling. The half-VG serial sections were analyzed (Total mice number is 5, half-VG has 5 segments, each segment has 3 sections). Data are means ± SD. \**P*<0.05 vs. *WT* control. **h,** Effect of Simvastatin on mouse Sca1^+^ cell growth. Growth curve of mouse Sca1+ cells cultured in stem cell growth medium with or without Simvastatin (0, 0.25, 0.5, 12.5μM) for 7 days. n=4, \**P* <0.05 vs. vehicle control, # *P* < 0.05 vs. 0.25 μM Sim, □ *P*< 0.05 vs. 0.25 and 0.5 μM Sim at each time point. **i,** Proliferation of mouse aortic SMCs cultured in different condition media (CMs), i.e., Ctl: stem cell basal media; Veh-CM: CM from Sca1^+^ VSCs without simvastatin (0.5μM); Sim-CM: CM from Sca1 ^+^VSCs with simvastatin (0.5 μM). n=4, \**P*<0.05 between the indicated groups. **j,** Migration and attachment of mouse aortic SMCs cultured in different CMs. **k,** Effect of Simvastatin on differentiation of mouse Sca1^+^ VSCs to endothelial cells. Representative pictures in the bright field and for the vWF and Sca1 immunofluorescence co-staining. **l,** The inhibition of peri-vascular simvastatin on VG remodeling at different time points. Representative immunofluorescence co-staining of Sca1/Ki67 on day 7 and Sca1/SMA on day 28. **m,** Upper panel, Quantified effect of local simvastatin on Sca1^+^ cell number on day 7 and 28. Three cross-sections were randomly chosen for each VG. n=6, \**P* <0.05 vs. control in the same group. Lower left panel, quantified proportions of double positive cells with Sca1 and Ki67 on day 7. Three cross-sections were randomly chosen for each VG. n=6, \**P*<0.05 vs. control in the same group. Lower right panel, quantified SMA positive area. 5 cross-sections of each VG were subjected to the analysis (n=6). \**P* < 0.05 vs. vehicle control. **n,** The inhibition of peri-vascular simvastatin on 100-day VG remodeling. Quantified SMA positive area of half-VG serial sections.5serial cross-sections of each VG were subjected to the analysis (n=4). The results are means ± SD. \**P*< 0.05 vs. vehicle control. **o,** Schematic illustration of Sca1^+^ VSC activation in VG remodeling. -, inhibition; + promotion.

In VGs, Sca1^+^ cells survive while SMCs (≈97%) and endothelial cells (ECs) (≈99%) die within 3 days after transplantation^1^ and the survived Sca1^+^ cells expand prior to the SMC proliferation leading to neointima formation^1^ (Figures 1a, 1b, S1, S2,and Tables S1, S2). Notably, the dynamics of intimal and adventitial Sca1^+^ cells in VG remodeling are different from that of Gli1^+^or CD34^+^cells (Figures 1a, 1b, S1, S2, and Tables S1, S2), suggesting that these cells may represent different subpopulations of VSCs in VGs. Indeed, co-staining of Sca1 with CD34, Gli1, SMA, or vWF revealed that Sca1^+^ cells represent a unique subset of VSCs and Sca1^+^CD34^+^ and Sca1^+^Gli1^+^ cells are subpopulations of Sca1^+^ VSCs in normal and remodeling veins (Figures 1a, 1b, S1, S2, and Tables S1 and S2). Collectively, these results suggest that Sca1^+^ VSCs may play a crucial role in VG intimal hyperplasia.

We generated *Sca1-Cre::Rosa26^floxedStop^eYfp* (*Sca1^Yfp^*) mice by crossing *Sca1-Cre* mice^3^ with *Rosa26^floxedStop^eYfp* reporter mice^1^ to genetically label Sca1^+^ cells with enhanced fluorescent yellow proteins (YFP) for tracking down the fate of Sca1^+^ cells in VG remodeling. The labeling efficiency was confirmed in various tissues including carotid arteries and jugular veins in 12-week-old *Sca1^Yfp^* mice (Figures S3, S4, S5, and Tables S3, S4). Intriguingly, there were a few YFP^+^Sca1^-^ SMCs in the tunica media of carotid arteries and jugular veins (Figure S5 and Tables S3, S4), revealing a novel subset of mature vascular SMCs derived from Sca1^+^ stem cells during development. We then carried out reciprocal transplantation of jugular veins between wild type (*WT*) and *Sca1^Yfp^* mice for 6 weeks (Figure 1c). We found that donor venous Sca1^+^ cells cannot survive, and only recipient Sca1^+^ cells repopulate the intima and the adventitia, but they do not contribute to neointima formation via SMC differentiation in VGs (Figures 1d, 1e, S6, S7, S8, S9, and Tables S3, S4). At proximal and distal anastomotic ends, but not the body, there are ≈2% of YFP^+^SMA^+^ cells in the neointima (Figures 1e, S6, and Table S3), reflecting the focal contribution of recipient arterial SMCs,^1^ in which some SMCs are labeled with YFP during the development (Figure S5 and Tables S3, S4). Notably, VG intimal Sca1^+^ VSCs are not likely derived from recipient intimal Sca1^+^ VSCs but other sources via the circulation (Figure S10). Since the Sca1^+^ cells isolated from aortic adventitia but not from the bone marrow could proliferate and have stemness in VSC growth media (data not shown), it is likely that VG adventitial Sca1^+^ VSCs are derived locally from the adjacent recipient arteries but not from the bone marrow.

We crossed *Sca1^Yfp^* mice with *Rosa26^floxedStop^DTR* (Cre-inducible transgenic expression of a diphtheria toxin receptor)^4^ mice to create *Sca1^Yfp^::Rosa26^floxedStop^DTR (iDTR)* mice, which render Sca1^+^ cells sensitive to an optimized regiment of diphtheria toxin (DT) treatment, thereby selectively depleting vascular Sca1^+^ cells in mice (Figures S11-12, and Table S6). We also noticed that the recipient Sca1^+^ cells are accumulated in the adventitia but not in the intima of VGs at 21 days after transplantation (Figure S12B). We then transplanted *WT* jugular veins into carotid arteries of control *WT* mice and vascular Sca1^+^ cell depleted *iDTR* mice for 21 days (Figure 1f). Since the VG anastomosis remodeling is complicated by YFP^+^SMCs derived from Sca1^+^ stem cells during development (Figures 1e, S6, and Table S4), we focused on the VG body in which neointimal cells are not derived from the recipient arterial SMCs^1^ or the donor venous Sca1^+^ cells (Figures 1e, S6, S8, and Table S4). Interestingly, the ablation of vascular Sca1^+^ cells in recipient mice significantly suppressed adventitial accumulation of Sca1^+^ cells and intimal hyperplasia in the VGs (Figure 1g), demonstrating a contributory role of the recipient derived adventitial Sca1^+^ cells to neointima formation in VGs. Because these Sca1^+^ cells reside in the adventitia and barely differentiate into SMCs in VGs (Figures 1e, S6, S8, and Tables S3, S4), it is likely that the adventitial Sca1^+^ cells promote intimal hyperplasia via paracrine regulation of the media SMC dedifferentiation towards synthetic SMCs in VGs.

Finally, we examined therapeutic potential of targeting adventitial Sca1^+^ VSCs in suppressing VG intimal hyperplasia using simvastatin, the clinically validated drug to treat intermediate and late VGF.^2^ We found that the therapeutic efficacy of simvastatin in suppressing VG intimal hyperplasia via a peri-vascular delivery for 3 days after transplantation is as effective as the oral administration 3 days prior to transplantation;^5^ however, simvastatin administration started 3 days after transplantation fails to inhibit the neointima formation (Figure S13). These unexpected results suggest that the primary target of simvastatin therapy is not likely vascular SMCs but VSCs that are the major cells survived within 3 days after transplantation (Figures 1a, 1b, S1, S2, and Table S1, S2). Indeed, simvastatin inhibited proliferation of aortic Sca1^+^ adventitial VSCs (Sca1^+^AdVSCs) (Figure 1h) but not aortic SMCs in vitro (Figure S14). It also suppressed Sca1^+^AdVSCs paracrine enforcement of medial SMC migration and proliferation (Figures 1i, 1j, S15). On the other hand, simvastatin promoted Sca1^+^AdVSCs differentiation to ECs and SMCs while inhibiting Sca1^+^AdVSCs differentiation to chondrocytes, adipocytes, and osteoblasts in vitro (Figures 1k, S16), revealing a simvastatin-induced naïve state of Sca1^+^ VSCs. Moreover, the peri-vascular simvastatin treatment almost blocked Sca1^+^AdVSCs proliferation associated with decreased accumulation of neointimal SMCs and suppressed intimal hyperplasia while increasing intimal Sca1^+^ cells in VGs (Figures 1l, 1m). These results suggest that simvastatin may convert recruited Sca1^+^ VSCs from an active form to a naïve state in VGs, thereby facilitating VG adaptation.

In summary, naïve Sca1^+^ VSCs are quiescent and geared for SMC and EC differentiation to maintain vascular homeostasis and repair. After transplantation, the recipient Sca1^+^ VSCs are activated and migrate to the adventitia and intima of VGs, where adventitial Sca1^+^ VSCs expand and gain the ability to paracrine enforce media SMC dedifferentiation while losing the potency for SMC or EC differentiation, and intimal Sca1^+^ VSCs lose the potential in EC differentiation (Figure 1o).

## DATA AVAILABILITY

The dataset used and/or analyzed during this study are available from the corresponding author on reasonable request.

## ACKNOWLEDEMENTS

This work as supported by grants from the National Institute of Biomedical Imaging and Bioengineering of the National Institute of Health (NIH) to T. Cui (R21EB022131) and from the National Heart, Lung, and Blood Institute of NIH to T. Cui (R01HL131667, R01 HL160541) and from the National Institute of General Medical Science of NIH to T. Cui (P20GM109091), and from the National Center for Complementary and Integrative Health of NIH to M. Nagarkatti, P. Nagarkatti and T. Cui (P01AT003961).

## AUTHOR CONTRIBUTIONS

W.W. and T.C. designed the experiments. W.W., H.Z., and L.Q. carried out the experiments, analyzed, and interpreted data. I.S-G. provided *Sca1-Cre* mice. M.C.M.W-E. provided *Rosa26^floxedStop^eYfp* mice. M.N., P.N., and T.C. provided research funds. W.W. and T.C. wrote, edited, and reviewed the manuscript. T.C. supervised the project. All authors have read and approved the manuscript.

## ADDITIONAL INFROMATION

### Supplementary information

Supplementary material is available online.

### Competing Interests

The authors declare no competing interests.

### Ethics declarations

All procedures in this study involving animals were reviewed and approved by the Institutional Animal Care and Use Committee for the University of South Carolina, USA.

## Notes

### Competing Interest Statement

The authors have declared no competing interest.

