## supplemental materials and figs for "Inactivation of Vascular Stem Cells Suppresses Intimal Hyperplasia in Vein Grafts"

**This word file includes:**

**Materials and Methods**

**Supplementary Text**

**Figures S1 to S17**

**Tables S1 to S5**

### Materials and Methods

#### 1. Material Table

| REAGENT or RESOURCE | SOURCE | IDENTIFIER |
|---|---|---|
| **Antibodies** | | |
| Mouse monoclonal anti-α-Smooth Muscle Actin (SMA) | Sigma Aldrich | Cat#: A5228 |
| Rat monoclonal anti-Mouse Ly-6A/E | BD | Cat#: 553333 |
| Rabbit monoclonal anti-Sca1 | Abcam | Cat#: ab109211 |
| Rabbit monoclonal anti-CD34 [EP373Y] | Abcam | Cat#: ab81289 |
| Rabbit polyclonal anti-GFP | Abcam | Cat#: ab6556 |
| Goat polyclonal anti-GFP | Abcam | Cat#: ab6673 |
| Rabbit polyclonal anti-Ki67 | Abcam | Cat#: ab15580 |
| Rat monoclonal anti-Gli1 | R&D | Cat#: MAB3324 |
| Rabbit polyclonal anti-vWF | Dako | Cat#: A0082 |
| Rabbit polyclonal anti-CD44 | Abcam | Cat#: ab18952 4 |
| Rat polyclonal anti-CD45 | BD | Cat#: 550539 |
| Mouse monoclonalanti-CD90 | Santa Cruz | Cat#: sc-53116 |
| Rabbit polyclonal anti-c-Kit | Abcam | Cat#: ab256345 |
| Mouse monoclonal anti-βactin | Sigma Aldrich | Cat#: a1978 |
| Mouse monoclonal anti- GAPDH | Sigma Aldrich | Cat#: G8795 |
| Alex fluor 488 | invitrogen | Cat#: A21208 |
| Alex fluor 488 | invitrogen | Cat#: A11001 |
| Alex fluor 488 | Invitrogen | Cat#: A21206 |
| Alex fluor 488 | Invitrogen | Cat#: A11055 |
| Alex fluor 546 | invitrogen | Cat#: A10036 |
| Alex fluor 546 | Invitrogen | Cat#: A10040 |
| Alex fluor 546 | Invitrogen | Cat#: A11081 |
| Alex fluor 546 | invitrogen | Cat#: A11056 |
| Anti-mouse IgG, HRP-linked antibody | Cell Signaling | Cat#: 7076 |
| Anti-rabbit IgG, HRP-linked antibody | Cell Signaling | Cat#: 7074 |
| **Chemicals** | | |
| Simvastatin | Sigma Aldrich | Cat#: S6196 |
| Diphtheria toxin | List biological laboratories, | Cat#: 15047A1 |
| leukemia inhibitory factor | Sigma Aldrich | Cat#: LIF2005 |
| 2-mercaptoethanol | Sigma Aldrich | Cat#: 63689-25ML-F |
| VEGF | Sigma Aldrich | Cat#: SRP6429 |
| 4',6-diamidino-2-phenylindole (Dapi) | Invitrogen | Cat#: D1306 |
| **Critical Commercial Assays** | | |
| In Situ Cell Death Detection Kit, TMR red | Roche | Cat#: 12156792910 |

| | | |
|---|---|---|
| StemPro® Adipogenesis Differentiation Kit | Gibco | Cat#: A10070-01 |
| StemPro® Chondrogenesis Differentiation Kit | Gibco | Cat#: A10071-01 |
| StemPro® Osteogenesis Differentiation Kit | Gibco | Cat#: A10072-01 |
| Human Endothelial SFM | Gibco | Cat#: 11111044 |
| Stem Cell Growth Medium | CellGenix® GMP | Cat#: 20802-0500 |
| Mouse ES Cell Basal Medium | ATCC® | Cat#: SCRR-2011 |
| **Software and Algorithms** | | |
| Volocity | PerkinElmer | |
| Image Pro Plus | MediumCybernetics | |
| Zeiss LSM Image Browser | Zeiss | |

### 2. Methods

#### Animals

Wild type (*WT*), *Rosa26*$^{floxedStop}$*eYfp* (*R26-eYfp*$^{fl/fl}$) and *Rosa26*$^{floxedStop}$*DTR* (the simian diphtheria toxin receptor) (*R26-DTR*$^{fl/fl}$) mice in a C57BL/6J genetic background were purchased from the Jackson Laboratory. *Sca1-Cre* mice in a C57BL/6J genetic background were kindly provided by Dr. Isidro Sánez-Garcia.[1] *Sca1-Cre::Rosa26*$^{floxedStop}$*eYfp* mice were generated by crossing *Sca1-Cre* mice with *Rosa26*$^{floxedStop}$*eYFP* mice. *Sca1-Cre::Rosa26*$^{floxedStop}$*eYfp:: Rosa26*$^{floxedStop}$*DTR* mice were generated by crossing *Sca1-Cre::Rosa26*$^{floxedStop}$*eYfp* mice with *Rosa26*$^{floxedStop}$*DTR* mice. Genotyping was performed by PCR analysis using mouse tail DNA (**Table S5**). Typical results of each genotype are shown in inserted figure.

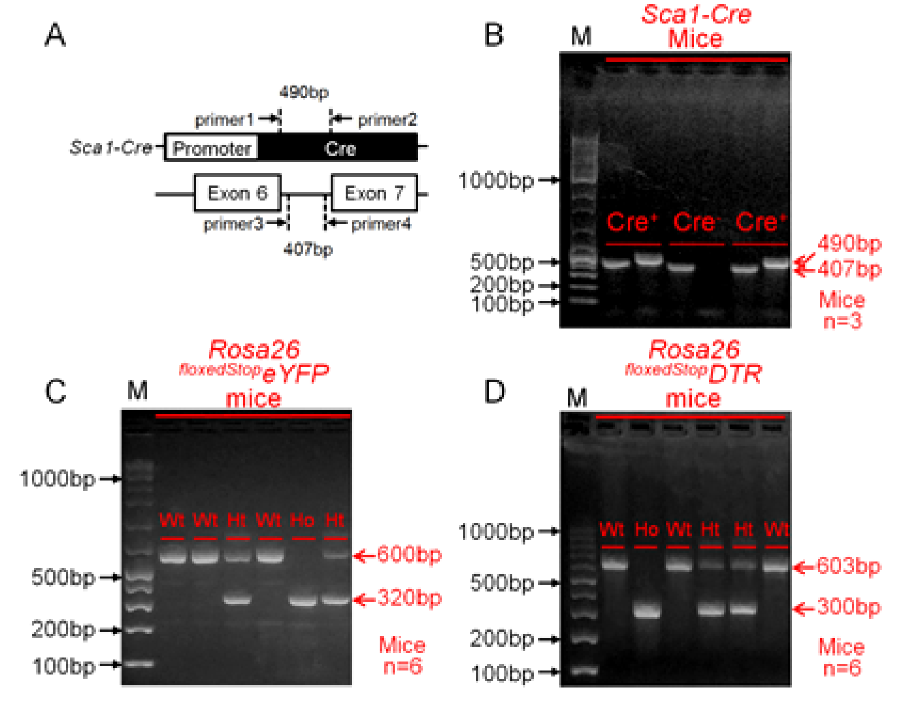

**Genotyping information. A**, Genomic locations of Sca1-Cre mice PCR products. **B-D**, Representative bands of PCR products in 1.5% agarose gel.

For the *Sca1-Cre* mice genotyping, the primer 1 & primer 2 amplified a 490 bp fragment which is the Cre allele (Cre: Bacteriophage P1 *Cre* gene for recombinase protein),[2-4] and primer 3 & primer 4 amplified a 407 bp fragment which is an internal positive control. Because the two PCR product bands were too close, so the primer 1 &

primer 2 and primer 3 & primer 4 were separated into 2 independent tubes with same PCR reaction system, then 2 gel electrophoresis lanes were used for each mouse. The genotyping of *R26-eYfp*$^{fl/fl}$ and *R26-DTR*$^{fl/fl}$ mice was carried out following the Jackson laboratory's protocols with minor modulations. The genotyping protocol(s) presented here (**Table S5**) have been optimized for reagents and conditions used by Dr. Cui lab. Only male mice were used in this study. All animals were housed at the AAALAC-accredited animal facility of University of South Carolina School of Medicine. All animals were treated in compliance with the USA National Institute of Health Guideline for Care and Use of Laboratory Animals. The use of animals and all animal procedures were approved by the Institutional Animal Care and Use Committee (IACUC) at University of South Carolina.

**Venous Bypass Graft Procedure**
Vein graft transplantation in mice was carried out as previously described.[5] Briefly, mice were anesthetized by intraperitoneal injection of a solution of xylazine (5 mg/kg body weight) with ketamine (80 mg/kg body weight) and fixed in supine position. Under a dissecting microscope (KL1500 LCD; Carl Zeiss, Jena, Germany), the right common carotid artery was mobilized and divided at the midpoint. Premade cuffs (Cat#:800/200/100/200, Portex Limited, United Kingdom) were placed on both ends of the artery, and the ends were reverted over the cuff and ligated. On the other hand, the right jugular veins of donor mice were exposed, excised and stored on ice in phosphate-buffered saline (PBS). The jugular vein segments were grafted between the 2 ends of the carotid artery by sleeving the ends of the vein over the artery cuff and ligating them together without changing the direction of blood flow.

**Simvastatin Administration**
Simvastatin (Cat#: S6196, Sigma-Aldrich) was delivered either locally or systemically as follows: Oral administration—Mice were treated with simvastatin at a dose of 1.6 mg/kg daily (1.6 mg/kg/d) by gavage from 3 days prior to the transplantation to 28 days after the transplantation, or from 3 days to 28 days after the transplantation. Perivascular administration—Simvastatin (30 µM) or phosphate-buffered saline in 50 µL 20% Pluronic F-127 gel (pH 7.2) (Cat#: P2443, Sigma-Aldrich) was applied to the adventitia of grafted vessels, immediately after the transplantation (**Figure S13**), which could maintain effective drug release between 1 and 3 days after the transplantation.[6]

**Tissue Harvest and Immunohistochemical Staining**
Organs including the brain, heart, kidney, spleen, liver, carotid artery, jugular vein, thoracic aorta, or vein grafts were harvested after the euthanasia with an overdose of pentobarbital or $CO_2$ inhalation and subject to immunohistochemical staining as we previously described.[5] The primary and secondary antibodies and other reagents used are listed in **Material Table.** Hematoxylin and eosin (HE) staining was carried out with Leica AutoStainer XL.

**Confocal Microscopy**
Confocal microscopic analysis was carried out using a Zeiss LSM510 META confocal scanning laser microscope and Volocity software (PerkinElmer, Waltham, MA, USA) as

we previously described.[5] A stained cross-section of vein grafts was scanned, and the image of a whole VG cross-section was integrated by combining individual image pieces with a black background using Power Point.

**Morphological Analyses of Vein Grafts**

Vein graft remodeling was assessed as we previously reported.[5] Briefly, the entire vein grafts was cut into serial 5-μm thick sections from proximal to distal ends, yielding around 900 sections (900 × 5 μm = 4.5 mm) per graft. These sections were uniformly divided into 9 segments with the number (#) 1 to 9, representing each 500 μm long segment from the proximal to distal end per graft. The area between inner edge of SMC positive area and inner edge facing the lumen was defined as the intimal area. A gap between adventitial inner edge (toward the lumen) and neointimal outer edge (toward the adventitia) was positioned, and the area between the gap and intimal inner edge and the area between the gap and adventitial outer edge were then defined as the neointimal area and the adventitial area, respectively. The circumferences) of the areas were measured by IPP (Image pro plus) software. We used the circumference to calculate the lumen and NI areas as well as the NI thickness and named it as C-method.

**Tracing Fate of Sca1$^+$ Cells in vivo**

The carotid arteries and jugular veins of male 12-week-old *Sca1-Cre::R26-eYfp$^{fl/-}$* report mice at physiological conditions (n=5) were harvested and processed for immunofluorescence staining of Sca1, YFP, and SMA. To quantify the labeling efficiencies in the vessels, 5 consecutive cross-sections with an interval of 100 μm were selected from each vessel for the analyses (**Figure S5**). The YFP-labeling efficiency of Sca1$^+$ cells in vivo was also determined by Western blot analysis and immunofluorescence staining of YFP protein expression in different organs (**Figures S3 and S4**).

At the experimental end point, the vein grafts were harvested and processed for immunofluorescence staining of YFP and SMA. The fate of YFP-labeled Sca1$^+$ cells in vein graft remodeling was quantified at 6 weeks after transplantation (**Figures S6, S7, S8 and S9**) as we previously described.[5]

**Ablation of Sca1$^+$ Cells in vivo**

*Sca1-Cre::R26-eYfp$^{fl/-}$::R26-DTR$^{fl/-}$* (*iDTR*) mice at age of 3 months were intraperitoneally (i.p.) injected with different doses of diphtheria toxin (DT) dissolved in PBS (List Biological Laboratories, USA). All male *iDTR* mice received i.p. injection of DT at a dose of 1.25 ng/g bodyweight/day (d), once every two days for 3 times, survived, whereas all male of female *iDTR* mice received one or two i.p. injection of DT at doses from 2.5 ng/g/d to 50 ng/g/d once every two days died (**Table S6**). Thus, male *iDTR* mice at age of 3 months received i.p. injections once every two days for 3 consecutive days followed by 3-day washout period were subject to vein graft transplantation surgery for 21 days (**Figure S11A**). Whilst the expression of YFP proteins in bone borrow of *iDTR* mice received the DT treatment was downregulated at the end of DT washout time (0 day) (**Figure S11B**), the number of YFP$^+$ cells was declined at 3 days and decreased dramatically at 21 days (Figure S11C). In addition, the numbers of

Sca1+ cells in the intima and adventitia were dramatically decreased at 3 days and undetectable at 21 days (**Figure S11D**). Accordingly, a regiment of DT treatment for ablating vascular Sca1+ cells in vivo is established. Parallelly, a set of experiments was designed to determine the impact of the *Cre*, *eYfp* and *DTR* gene insertion on vein graft remodeling. Jugular veins of male *WT* mice at age of 3 months were transplanted into carotid arteries of 3-month-old male *WT* mice and *iDTR* mice without the DT treatment for 21 days (n=4-5) (**Figure S12A**). In the vein grafts transplanted into the *iDTR* mice, there was a substantial amount of Sca1+ cells accumulated in the adventitia but barely detected in the intima (**Figure S12B**). The sizes of neointima in vein grafts transplanted to *WT* and *iDTR* mice were comparable (**Figure S12B-C**). These results demonstrate a minimal effect of the gene manipulation for ablating vascular Sca1+ cells per se on vein graft remodeling at least under our experimental setting. Finally, jugular veins of male *WT* mice at age of 3 months were transplanted to carotid arteries of male *iDTR* mice at age of 3 months after receiving the optimized regiment of DT treatment for 21 days (n=5).

**Cell Culture**

**i. Mouse Aortic Smooth Muscle Cells** were isolated and cultured as we previously described.[5] Briefly, 5 thoracic aorta segments (the length of each segment is ~1-cm) of 2-3 –month-old male wild type mice in a C57BL/6J genetic background were removed, pooled, and subject to enzymatic digestion using type II collagenase (2 mg/ml; Cat#: 4177, Worthington Biochemical Corp., USA), soybean trypsin inhibitor (1 mg/ml, Cat#: LS003571, Worthington Biochemical Corp., USA) and elastase III (0.744 units/ml; Cat#: LS002279, Worthington Biochemical Corp., USA) for 1-2 h at 37°C with periodically shaking throughout the incubation. The digestion was terminated by addition of DMEM containing 20% FBS (Cat#: FBS-500, X& Cell Culture, USA). Finally, the enzymatically isolated single cells (~7000/per aorta, ~35000/~5 aortas) were cultured in 1 well of a 6-well plate with high glucose (4.5 g/L) DMEM (Cat#: 11995, Gibco, USA) supplemented with 20% FBS and 100 µg/ml penicillin/streptomycin (Cat#: 30-002-CI, Corning, USA) at 37°C in a 5% $CO_2$ incubator. The cultured mSMCs between passage (P)3 and P10 in which ~99% of the cells were stained positively for αSMA were used in this study.

**ii. Mouse Aortic Adventitial Sca1+Cells** were isolated and cultured as previously described.[2] Briefly, 2~3-month-old male mice (n=5) were euthanized by $CO_2$ overdose inhalation and then the ascending aortas, aortic arches, thoracic aortas (the length of each thoracic aorta is ~2-cm) were removed, washed in phosphate-buffered saline (PBS) and incubated in 4.5 g/L glucose Dulbecco's modified Eagle's medium (DMEM) containing 1 mg/ml collagenase type II (Cat#: 17101-015, Worthington Biochemical Corp.) at 37°C for 10-15 min. The surround connective tissue and the adventitia were separated from the aortic medium. The dissected adventitia were chopped into small pieces (~1 $mm^2$). The adventitia pieces were digested in 2.5 ml of type II collagenase (14 mg/ml; Cat#: 4177, Worthington Biochemical Corp., USA) for 1-2 hours, and then cold 20% FBS-DMEM were used to terminate the digestion. Sca1+ adventitial cells were isolated using Anti-Sca1 MicroBead Kit (MiltenyiBiotec GmbH, BergischGladbach, Germany, order no.130-092-529) according to the manufacturer's instructions. Briefly, the enzymatically isolated adventitial cells were washed with PBS containing 0.5% BSA, and incubated with anti–Sca1 immunomagnetic microbeads. The cell suspension was added to a column equipped with a magnetic cell sorting system (MACS). After washing, Sca1$^+$ cells were collected. For higher purity of Sca1$^+$ cells, a second column was used to repeat the process. About 0.5~1 × $10^4$ cells were seeded in a well of a 6-well plate for 3 days with stem cell growth medium (SCGM) (CellGenix® GMP Stem Cell Growth Medium) supplemented with10 ng/ml leukemia inhibitory factor and 10% FBS) without touching. 3 days later, changed the medium every 2 days. The Sca1$^+$cells up to 10 passages were used in this study.

**iii. Mouse Bone marrow Sca1$^+$Cells** were isolated and cultured as previously described.[7] Briefly, one male wild type C57BL/6J mice at age of 2-3 months were used for one isolation. Bone marrow cells were flushed out from the limb. Sca1$^+$ cells in the total bone marrow cells were isolated by microbeads according to the manufacturer's instructions. Briefly, cells were washed with PBS containing 0.5% BSA, and incubated with anti–Sca1 immunomagnetic microbeads (MiltenyiBiotec GmbH, BergischGladbach, Germany). The cell suspension was added to a column equipped with a magnetic cell sorting system (MACS). After washing, Sca1$^+$ cells were collected. For higher purity of Sca1$^+$ cells, a second column was used to repeat the process. Bone marrow cells were cultured with BM medium (DMEM medium with 15% FBS). Then change medium 16 h later after seeding. After that, the medium was changed every other day.

**Proliferation of Mouse Aortic Adventitial Sca1$^+$Cells**

Mouse aortic adventitial Sca1$^+$ cells at passage 6 were seeded into 12-well cell culture plates (1.5 x $10^4$ per well) in SCGM (CellGenix® GMP Stem Cell Growth Medium) supplemented with10 ng/ml leukemia inhibitory factor and 10%FBS) with or without simvastatin (0, 0.25, 0.5, 1 μM) for 7days. The culture medium was changed on day 3 and the cell numbers per well was counted on day 1, day 3 and day 5 after the seeding (**Figure 1h**).

**Preparation of Conditioned Medium (CM) of Mouse Aortic Adventitial Sca1$^+$Cells**

Sca1$^+$ cells at a 50% of confluent state were cultured in SCGM growth medium with or without simvastatin (0.5 μM) for 3 or 7 days. These cells were washed three times with PBS, and the culture medium were replaced with SCGM free medium and incubated for 24 h. Thereafter, the culture medium was collected, and filtered by 2.2μM filter (Cat#: 6794-2502, Whatman, UK), and then stored in -80°C freezer (**Figures S15 and S16**). The conditional medium (CM) from Sca1$^+$ cells without treatment of simvastatin (0.5 μM) was designated as Veh-CM and conditional medium from Sca1$^+$ cells in response to simvastatin (0.5 μM) treatment were named as Sim-CM, respectively (**Figures S15 and S16**).

**Assessments of mouse Vascular SMC Dedifferentiation**

**i. Proliferation Assay**—Mouse aortic SMCs at passage 10 were seeded into 12-well cell culture plates (1.5 x $10^4$ per well) in the different Sca1$^+$ cells-conditioned media supplemented with 1% FBS (Cat#: FBS-500, X & Cell Culture, USA). The culture medium was changed on day 3 and the cell numbers per well was counted on day 1, day 3 and day 5 after the seeding (**Figures 1i, S14 and S15D**).

**ii. Migration Assay**—Mouse aortic SMCs at passage 10 were seeded into 12-well cell culture plates (1.5 x 104 per well) in SMC growth medium, i.e., high glucose DMEM (Cat# 11-965-092, Gibco, USA) supplemented with 10% FBS (Cat#: FBS-500, X& Cell Culture, USA) for 48 h, reaching a confluent state. After serum-starvation for 24 h, a wound on the confluent SMC layer was created by a 200 µl tip and then cultured with SCGM free medium (Ctl-SCGM), vehicle (Veh)-treated $Sca1^+$ cells conditioned medium (Veh-CM), Sen A-CM or Sen B-CM with 0.1% FBS for 24 h and 48h. The wounds were captured by a microscope (Nikon E600 Widefield Epifluorescence and Darkfield Microscopy, Nikon, Japan) on day 0, day 1 and day 2 after the scratch. The migration area was measured using IPP (Image Pro Plus) analysis software (**Figures 1j and S15**).

**iii. Attachment Assay** —Mouse aortic SMCs at passage 10 were seeded into 12-well cell culture plates (1.5 x $10^4$ per well) with SCGM free medium (Ctl-SCGM), vehicle (Veh)-treated $Sca1^+$ cells conditioned medium (Veh-CM), Sen A-CM or Sen-B CM without any supplements for 24 h, then got pictures by a microscope (Nikon E600 Widefield Epifluorescence and Darkfield Microscopy, Nikon, Japan) on day 1 after seeding (**Figure 1j**).

**Assessments of mouse $Sca1^+$ Cells Differentiation**

Mouse $Sca1^+$ cells at passage 8 were cultured in SCGM growth media. When the confluency reached to 100%, changed to media for different differentiation media. For the adipocyte, StemPro® Adipogenesis Differentiation Kit (A10070-01) and Oil Red O staining were used. For chondrocyte, StemPro® Chondrogenesis Differentiation Kit (A10071-01) and Alcian Blue staining were used. For osteogenesis, StemPro® Osteogenesis Differentiation Kit (A10072-01) and Alizarin Red staining were used. For smooth muscle cells, DMEM (GIBICO-10569) + 10%FBS medium and SMA/CNN1 immunofluorescence staining were used. For endothelial cells, Human Endothelial SFM (GIBICO) + 10%FBS + 100 µg/ml penicillin/streptomycin + 10 ng/mL VEGF medium and vWF immunofluorescence staining were used. Bright field pictures were taken on day 3 and the last day during differentiation (**Figure S16**).

**Polymerase Chain Reaction (PCR)**

Genomic DNAs extracted from mouse tails were subject to PCR for genotyping of transgene mice using the primers and reaction conditions as shown in **Table S5**.

**Western Blot Analysis**

Snap-frozen tissues were smashed into powder in liquid nitrogen and then lysed in a homogenization buffer (RIPA) contained 150 mM NaCl, 1% NP-40, 50 mM Tris (pH 8.0), 0.5% sodium deoxycholate (C24H39NaO4), 0.1% sodium dodecyl sulfate (SDS), and proteinase and phosphatase inhibitors (Cat#: P8340 and P0044, Sigma-Aldrich, USA), and subject to Western blot analysis as we previously described.[5] The primary and secondary antibodies used are listed in **Material Table**.

**Statistical analysis**
Data are shown as mean ± standard error of the mean (SEM). Differences between 2 groups were evaluated for statistical significance with the Student's t-test, and differences were considered significant at $P < 0.05$.

### Supplementary Text

#### 1. Cellular Dynamics of Vein Graft (VG) Remodeling

All mice used for vein graft (VG) transplantation were male 3-month-old wild type mice in a C57BL/6J genetic background; and vessels were harvested at different time (0, 1, 3, 7, 14, 42 days) after transplantation. Vessel cross-sections were subject to morphological analyses of endothelial cells (ECs) identified by vWF (Von Willebrand factor) staining, SMCs by SMA staining, VSCs by Sca1, Gli1, and CD34 staining, apoptosis by Tunel staining, and proliferation by Ki67 staining.

In normal murine jugular veins, most of the $Sca1^{+}$cells resided mainly in the intima and the adventitia, making up approximately ≈60% of intimal cells and ≈40% of adventitial cells, respectively (**Figure S1, and Tables S1**). Similarly, most of the $CD34^{+}$ cells were in the intima and the adventitia, contributing to ≈70% of intimal cells and ≈80% of adventitial cells, respectively (**Figure S1 and Table S1**). However, $Gli1^{+}$ cells were located predominantly in the adventitia, constituting ≈80% of adventitial cells (**Figure S1 and Table S1**). In addition, there are 43% of intimal cells double positive for Sca1 and vWF and 44% double positive for Sca1 and CD34 (**Figure S2**), indicating the heterogeneity of intimal $Sca1^{+}$ cells. Similarly, the heterogeneity of adventitial $Sca1^{+}$ cells is evidenced by ≈50% of $Sca1^{+}CD34^{+}$ and ≈30% of $Sca1^{+}Gli1^{+}$ adventitial cells (**Figure S2 and Table S2**).

In VGs, intimal $Sca1^{+}$ cells disappeared at 1 day and were hardly observed until 14 days but completely recovered at 42 days; whereas the number of adventitial $Sca1^{+}$cells was decreased at 1 day but then was quickly recovered at 3 days (≈20%) and kept increasing until 42 days (≈70%) (**Figures S1, S2, and Tables S1, S2**). Also, few scattered $Sca1^{+}$ cells were found in the neointima from day 7 to day 42 (**Figures S1, S2, and Tables S1, S2**). Like the adventitial $Sca1^{+}$ cells, the number of adventitial $Gli1^{+}$ cells was decreased at 1 day, and then, however, unlike the $Sca1^{+}$ cells, the adventitial $Gli1^{+}$ cells were not recovered until 42 days (≈80%) (**Figures S1, S2, and Table S1**). All $CD34^{+}$ intimal cells disappeared at 1 day and were barely detected until the reendothelialization at 42 days (**Figures S1, S2, and Table S1**). The adventitial $CD34^{+}$ cells exhibited a similar pattern to adventitial $Sca1^{+}$ cells up to 42 days; however, their numbers were less than that of $Sca1^{+}$ cells (**Figures S1, S2, and Table S1**). The $CD34^{+}$ neointimal cells exhibited a similar pattern to the $Sca1^{+}$ cells (**Figures S1, S2, and Tables S1, S2**). These results suggest that these cells may represent different subpopulations of VSCs in VGs. Indeed, co-staining of Sca1 with CD34, Gli1, SMA, or vWF revealed that in normal jugular veins at baseline, the of intimal or adventitial numbers $Sca1^{+}CD34^{+}$ cells are like that of $Sca1^{+}$cells but are less than that of $CD34^{+}$cells (**Figures S1, S2, and Table S1, S2**). However, in remodeling VGs up to 42 days after transplantation, the numbers of intimal $Sca1^{+}CD34^{+}$ cells are similar to that of intimal Sca1$^+$ or CD34$^+$ cells, and the numbers of adventitial Sca1$^+$CD34$^+$ cells are much less than that of adventitial Sca1$^+$or CD34$^+$cells (**Figures S1, S2, and Table S1, S2**). In addition, adventitial numbers of Sca1$^+$Gli$^+$ cells are less than that of Sca1$^+$ or Gli1$^+$ cells in both normal jugular veins at baseline and remodeled VGs at 42 days after transplantation, while they are hardly observed between day 1 to day 14 (**Figures S1, S2, and Tables S1, S2**). These results suggest that Sca1$^+$ cells represent a unique subset of VSCs and Sca1$^+$CD34$^+$ and Sca1$^+$Gli1$^+$ cells are subpopulations of Sca1$^+$ VSCs in normal and remodeling veins.

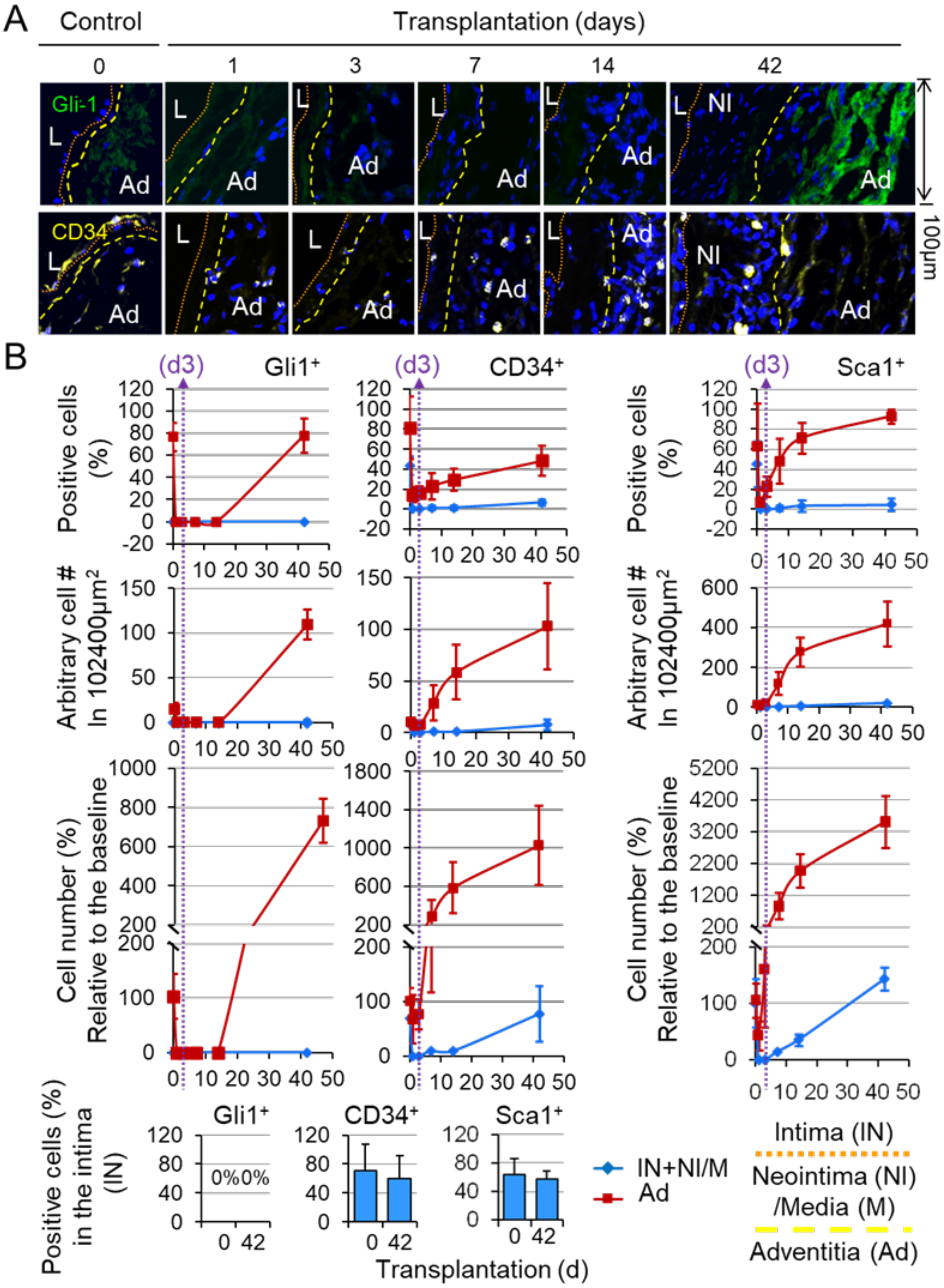


**Figure S1. Vascular Stem Cell (VSC) Dynamics in Vein Graft Remodeling.** Vein graft (VG) transplantation was carried out using male 3-month-old WT mice in a C57BL/6J genetic background. VGs were harvested at 0, 1, 3, 7, 14, 42 days after transplantation and subject to morphological analyses as described in Methods (n=8). (**A**) Representative immunofluorescence staining for Gli1 (green), CD34 (yellow). Small dotted red lines separate IN and NI&M; big dotted yellow lines separate NI&M and Ad (IN, the intima; NI, the neointima; M, the media; Ad, the adventitia). (**B**) Quantified percentages and numbers of the cells positive for VSC markers as indicated in 2 different layers (IN+NI&M and Ad). Four cross-sections were randomly chosen at each time point per VG (n=8) and subject to the immunofluorescence staining analysis. All the pictures which could be used for analysis were counted. Percentages of positively stained cell numbers as well as the arbitrary numbers for positively stained cells in a 102400μm$^2$ area in each indicated layer are quantified. Percentages of the cell numbers relative to the mean number in normal vein in each indicated layers are also quantified (blue lines indicate IN+NI&M; red lines indicate Ad).

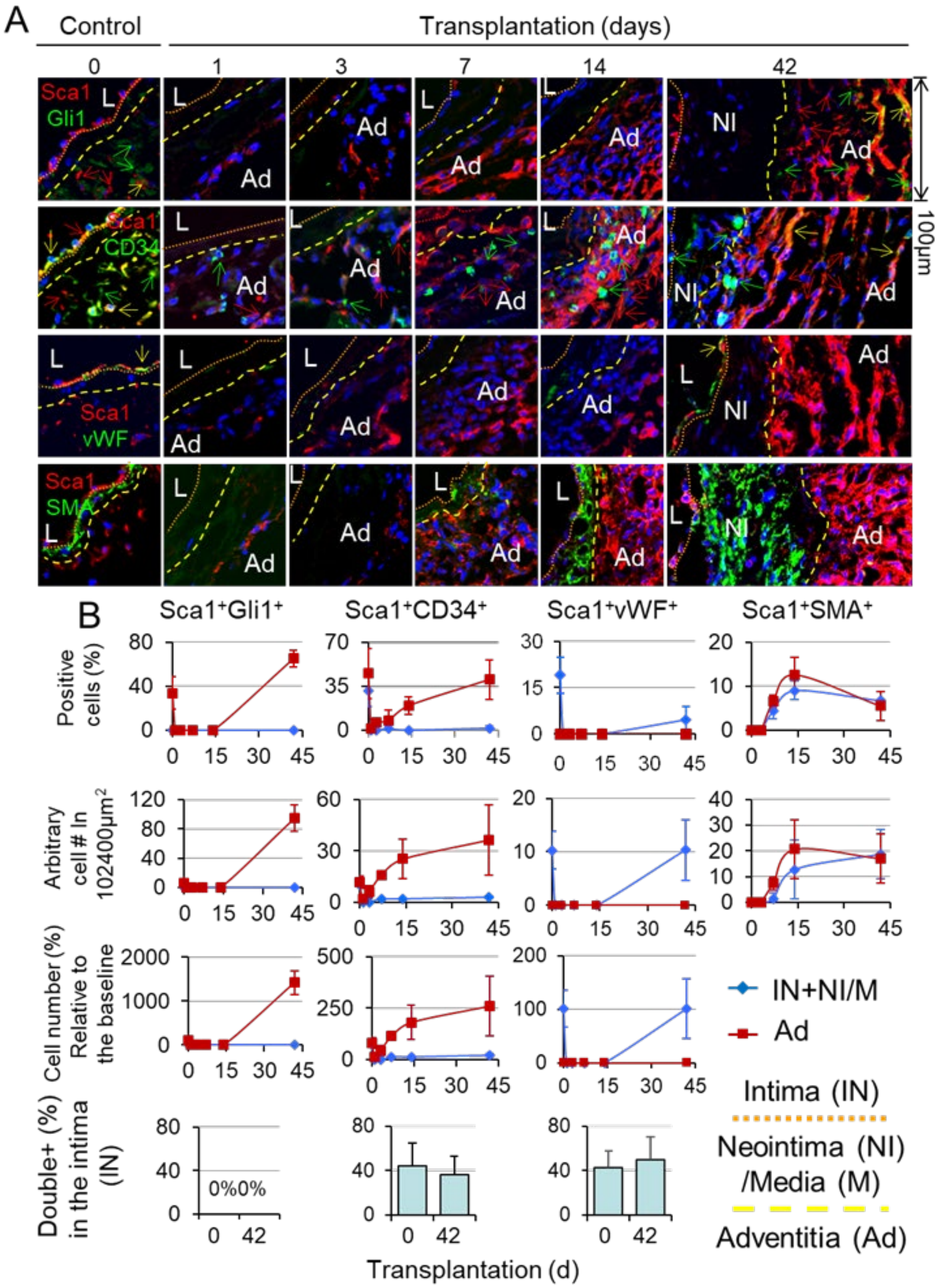


**Figure S2. The heterogeneity of Vascular Stem cells (VSCs) in Vein Graft Remodeling.** Vein graft transplantation, vessel tissue harvest, and morphological analysis were carried out as in **Figure S1** (n=8). (**A**) Representative immunofluorescence co-staining of Sca1(red) with vWF, SMA, CD34, Gli1 (they are all green) in VGs (Control, normal jugular vein before surgery 0 day and VGs at 1, 3, 7, 14, 42 days after transplantation were used for time course study) (IN, the intima; NI, the neointima; M, the media; Ad, the adventitia). Small dotted red lines separate IN and NI&M; big dotted yellow lines separate NI&M and Ad. (**B**) Quantified percentages and numbers of the cells double positive for VSC markers as indicated in 2 different layers (IN+NI&M and Ad). Four cross-sections were randomly chosen at each time point per VG (n=8) and subjected to the immunofluorescence staining analysis. All the pictures which could be used for analysis were counted. Percentage of positively stained cell numbers as well as the arbitrary numbers of positively stained cells in a 102400$\mu m^2$ area in each indicated layer is quantified. Percentages of the cell numbers relative to the mean number in normal vein in each indicated layers are also quantified (blue line indicates IN+M&NI; red line indicates Ad).

**2. The Fate of Sca1$^+$ Cells in Vein Graft Remodeling**

*Sca1-Cre::Rosa26$^{floxedStop}$eYfp* (*Sca1$^{Yfp}$*) mice were generated by crossing *Sca1-Cre* mice[1] with *Rosa26$^{floxedStop}$eYfp* reporter mice[5] to genetically label Sca1$^+$ cells with enhanced fluorescent yellow proteins (YFP) for tracking down the fate of Sca1$^+$ cells in VG remodeling. Baseline characterization revealed that Sca1$^+$ hematopoietic cells, liver non-hepatic cells, kidney cortical tubular cells, cardiac interstitial cells, and vascular endothelial cells are effectively labeled by YFP in 12-week-old mice as previously reported[8,9] (**Figures S3, S4**), demonstrating the genetic labeling efficacy. There are ≈54% of intimal cells and ≈49% of adventitial cells labeled with YFP in the carotid artery (**Figure S5A and Table S3**). Also ≈76% of intimal cells and ≈51% of adventitial cells labeled with YFP in jugular veins (**Figure S5B and Table S4**). Intriguingly, there are ≈9% and ≈2% of YFP$^+$Sca1$^-$ SMCs in the tunica media of carotid arteries and jugular veins, respectively (**Figure S5 and Tables S3-4**), revealing a novel subset of mature vascular SMCs derived from Sca1$^+$ stem cells during development.

Next, reciprocal transplantation of jugular veins was carried out between wild type (*WT*) and *Sca1-Cre::Rosa26$^{floxedStop}$eYfp* (*Sca1$^{Yfp}$*) mice for 6 weeks. Each VG from the proximal to the distal end was sectioned and analyzed as we recently reported.[5] In the *WT* VGs transplanted into the carotid arteries of *Sca1$^{Yfp}$* mice, ≈40% of intimal cells, ≈10% of neointimal cells and ≈80% of adventitial cells were positive for YFP across entire grafts (**Figures S6, S7, and Table S3**). None of YFP$^+$ cells in the intima were positive for SMA (Figure S6). At proximal and distal anastomotic ends, but not the body, there were ≈2% and ≈8% of YFP$^+$SMA$^+$ cells in the neointima and the adventitia, respectively (**Figures S6, S7, and Table S3**). In sharp contrast, no YFP signal could be detected in any anatomic locations of the *Sca1$^{Yfp}$* VGs transplanted into *WT* mice (**Figures S8, S9, and Table S4**). In addition, intimal number of YPF$^+$ cells was larger than that of Sca1$^+$ cells in normal carotid arteries (CAs) of *Sca1$^{Yfp}$*mice (**Figure S10**), suggesting that some endothelial cells are derived from the intimal Sca1$^+$VSCs and intimal number of YFP$^+$Sca1$^-$ cells reflects the intimal Sca1$^+$ VSCs differentiation into endothelial cells. There were less intimal YFP$^+$Sca1$^-$ cells in the *WT* VGs transplanted into *Sca1$^{Yfp}$* mice compared to normal carotid arteries of *Sca1$^{Yfp}$* mice (7% vs. 30%), whereas there is no difference of intimal YFP$^+$ cells between the VGs and carotid arteries of *Sca1*$^{Yfp}$ mice (**Figure S10**). These results not only validate that recipient arterial endothelial cells are the source of reendothelialization of VGs,[5] but also reveal that the stemness of intimal Sca1$^{+}$ VSCs towards endothelial cell differentiation in VGs is reduced, suggesting that VG intimal Sca1$^{+}$ VSCs are not likely derived from recipient intimal Sca1$^{+}$ VSCs but other sources via the circulation.

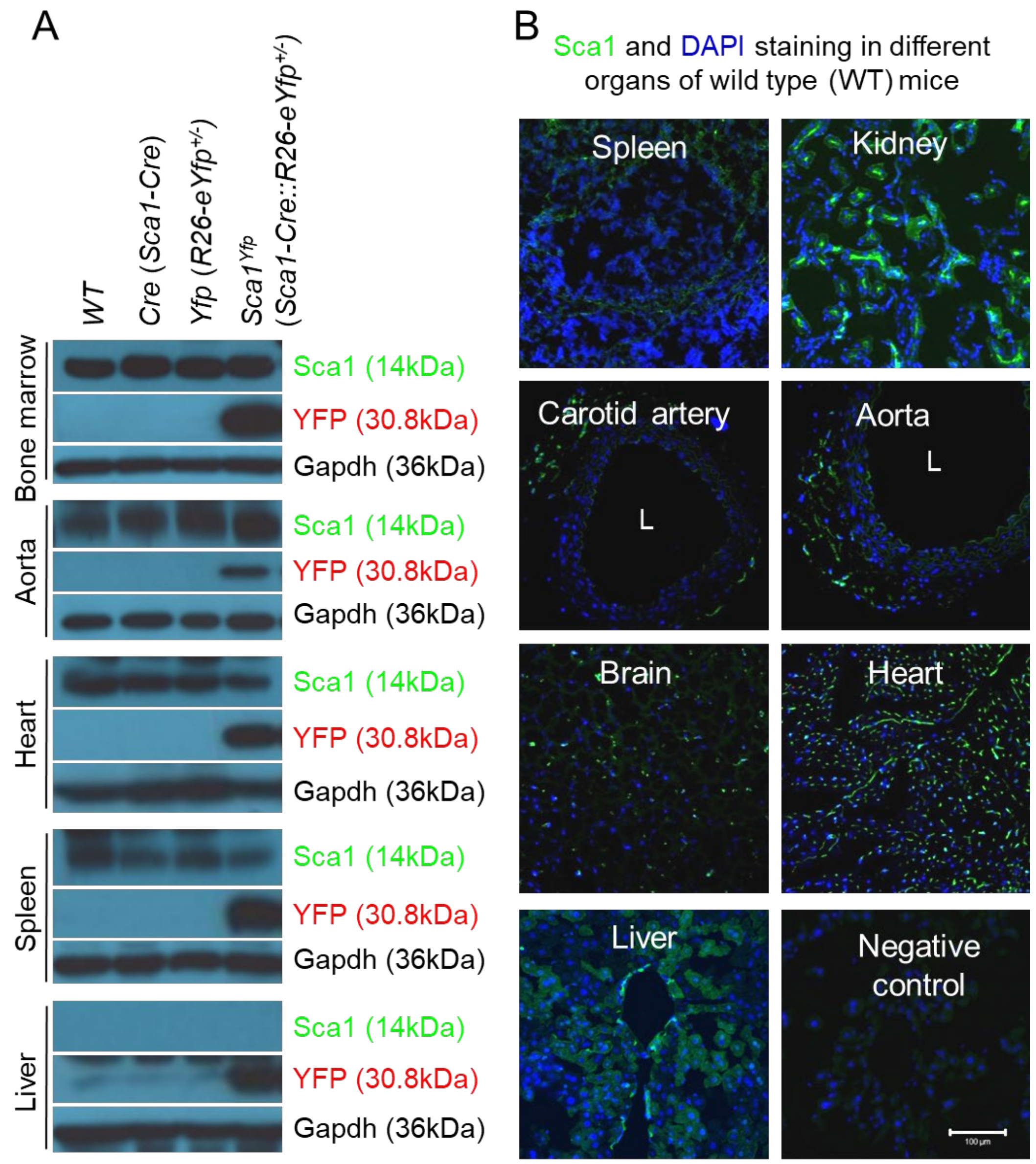


**Figure S3.** Baseline Characterization of *Sca1-Cre::Rosa26*$^{floxedStop}$*eYfp* mice. Male 12-week-old wild type (*WT*), *Sca1-Cre* (*Cre*), heterozygotes of *Rosa26*$^{floxedStop}$*eYfp* (*R26-eYfp*$^{fl/-}$), and heterozygotes of *Sca1-Cre::Rosa26*$^{floxedStop}$*eYfp* (*Sca1-Cre::R26-eYfp*$^{fl/-}$; *Sca1*$^{Yfp}$) mice (n=5) were used for Western blot analysis and immunofluorescence staining of YFP protein expression in different organs. (**A**) Western blot analysis of Sca1, YFP and Gapdh expression in different organs including the bone marrow, aorta,

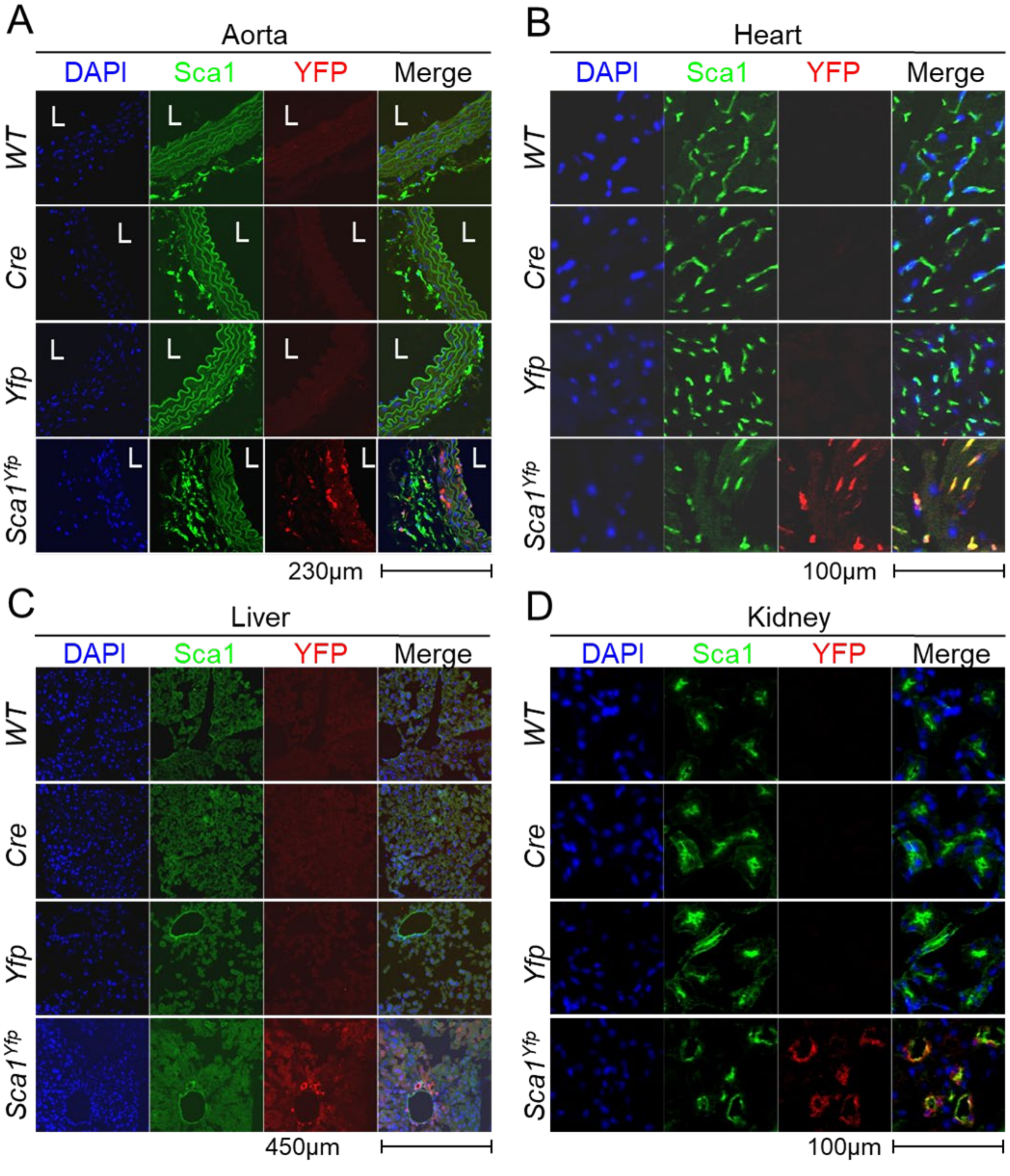


heart, spleen, and liver. (**B**) The representative immunofluorescence Sca1 staining in different organs including the spleen, kidney, brain, heart, carotid artery, aorta, and liver.

**Figure S4. Baseline Characterization of *Sca1-Cre::Rosa26$^{floxedStop}$eYfp*mice**.Male 12-week-old wild type (*WT*), *Sca1-Cre* (*Cre*), heterozygotes of *Rosa26$^{floxedStop}$eYfp* (*R26-eYfp$^{fl/-}$*), and heterozygotes of *Sca1-Cre::Rosa26$^{floxedStop}$eYfp* (*Sca1-Cre::R26-eYfp$^{fl/-}$*; *Sca1$^{Yfp}$*) mice (n=5) were used for immunofluorescence staining of YFP and Sca1protein expression in (**A**) the aorta, (**B**) the heart, (**C**) the liver, and (**D**) the kidney..

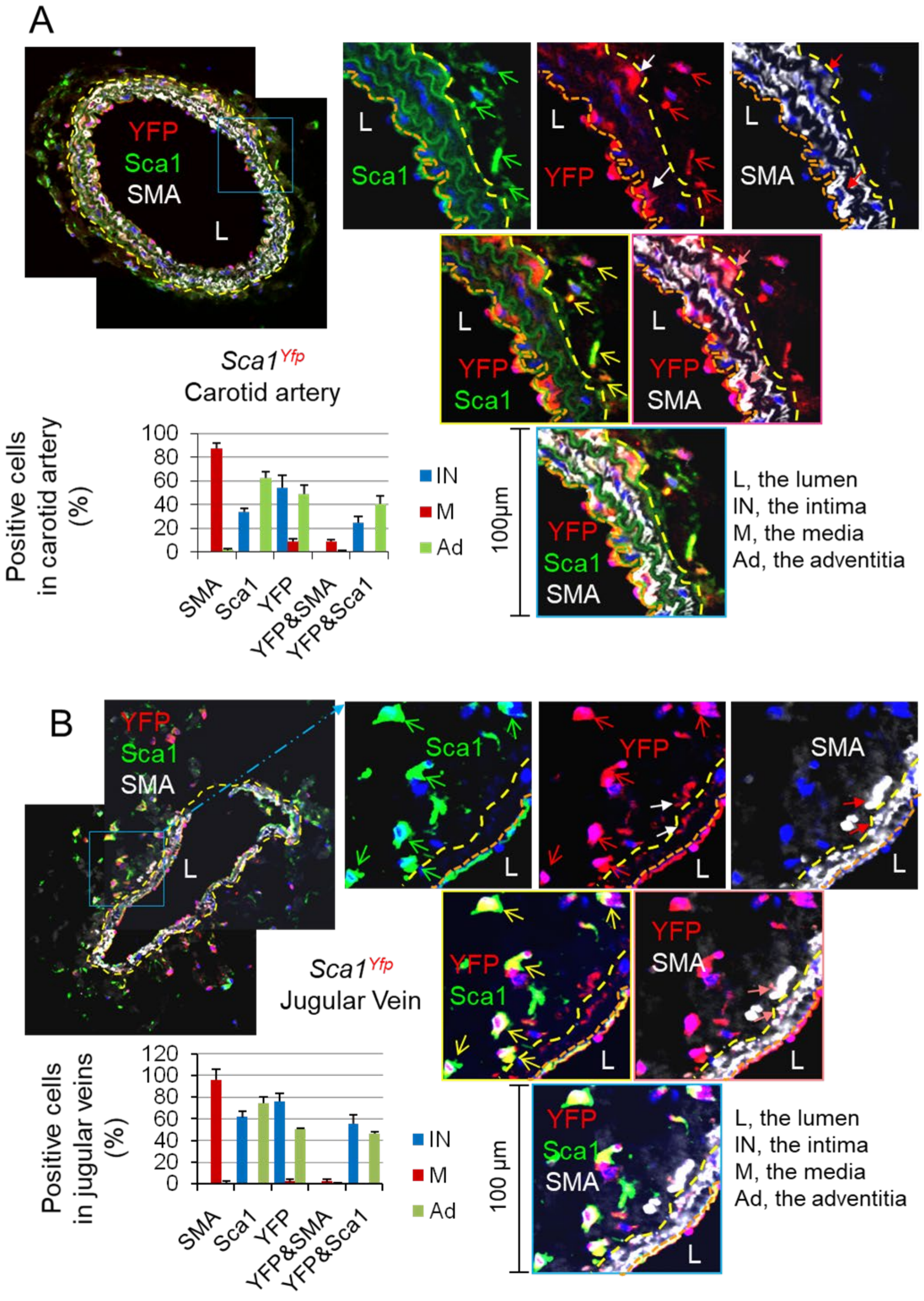


**Figure S5. Baseline Characterization of *Sca1-Cre::Rosa26$^{floxedStop}$eYfp* mice.** Male 12-week-old wild type (*WT*) and heterozygotes of *Sca1-Cre::Rosa26$^{floxedStop}$eYfp* (*Sca1-*

*Cre::R26-eYfp*$^{fl/-}$; *Sca1*$^{Yfp}$) mice(n=3) were used for immunofluorescence staining of YFP, Sca1, and SMA protein expression in the carotid artery and jugular vein. *WT* mice were used as control. The representative images of YFP, Sca1, and SMA staining on whole vessel cross-sections of normal jugular veins (**A**) and carotid arteries (**B**) in *Sca1*$^{Yfp}$ mice. YFP is red; SMA is white, Sca1 is green. L, the lumen; IN, the intima; NI, the neointima; Ad, the adventitia. Dotted orange lines separate the IN and NI and yellow lines separate the NI and Ad layers. The percentages of cells positive for different markers in each major layer of jugular veins and carotid arteries were quantified. Data were showed as means ± SD (5 whole vessel cross-sections per mice, 15 whole vessel cross-sections were analyzed).

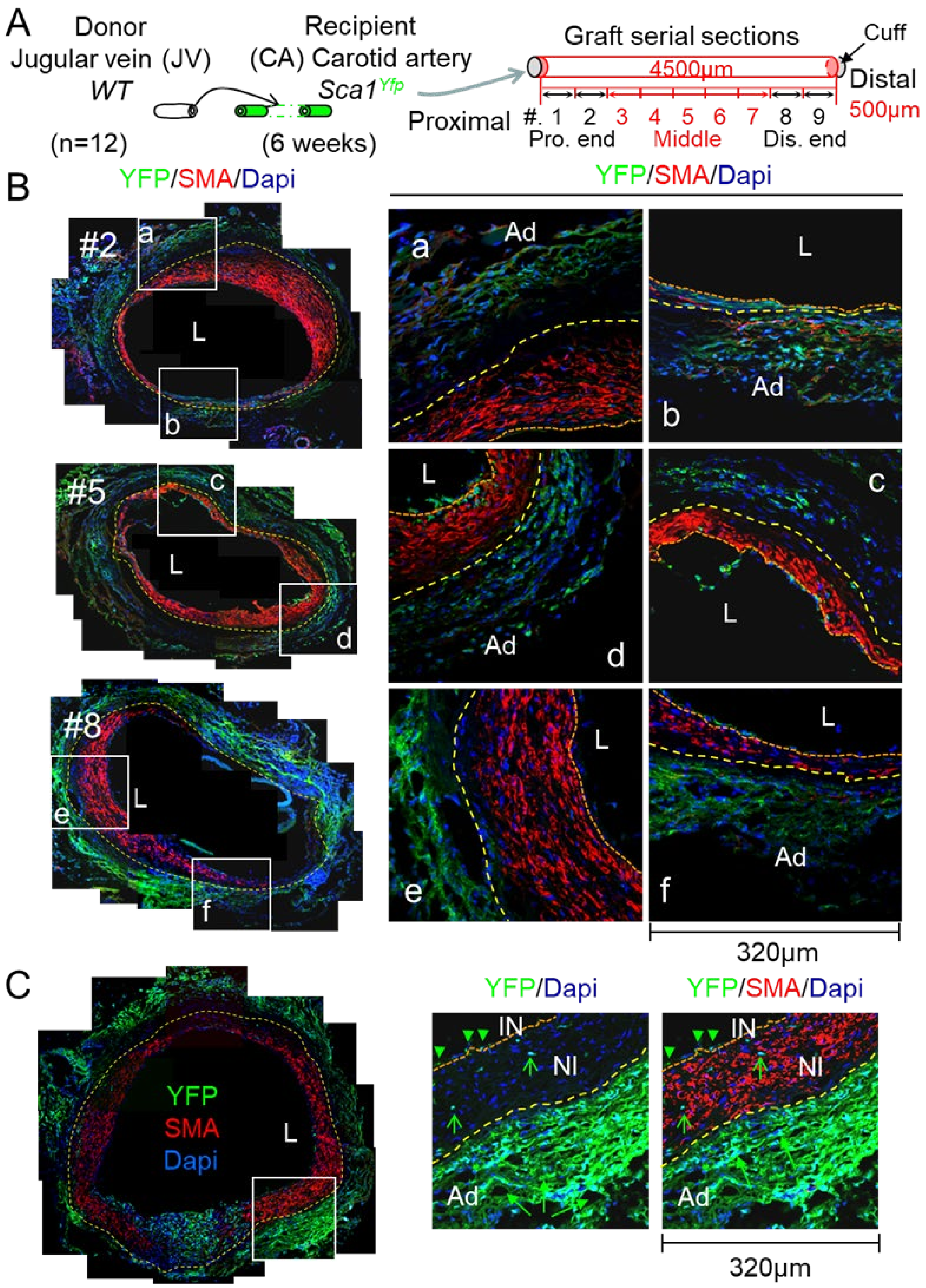


**Figure S6. The Contribution of Recipient Sca1⁺ Cells to Neointima Formation in VGs.** (**A**) A Scheme of VG transplantation and cross-tissue sectioning. **Pro.end**, proximal anastomotic end (#1-2); **Middle**, middle body (#3-7); **Dis.end**, distal anastomotic end (#8-9). (**B**) The representative images of co-staining of YFP with SMA in whole cross-sections of each segment (#) of wild type (*WT*) jugular vein (JV) grafts transplanted to carotid arteries (CAs) of *Sca1*$^{Yfp}$ mice in which Sca1$^+$ cells were genetically labeled with YFP (n=12). (**C**) The representative images of co-staining of YFP with Sca1 in whole cross-sections of *WT*-JV grafts transplanted to CAs of *Sca1*$^{Yfp}$ mice for **Figure 1e**. YFP is green; SMA is red. L, the lumen; IN, the intima; NI, the neointima; Ad, the adventitia. Dotted orange lines separate the IN and NI, and yellow lines separate the NI and Ad layers.

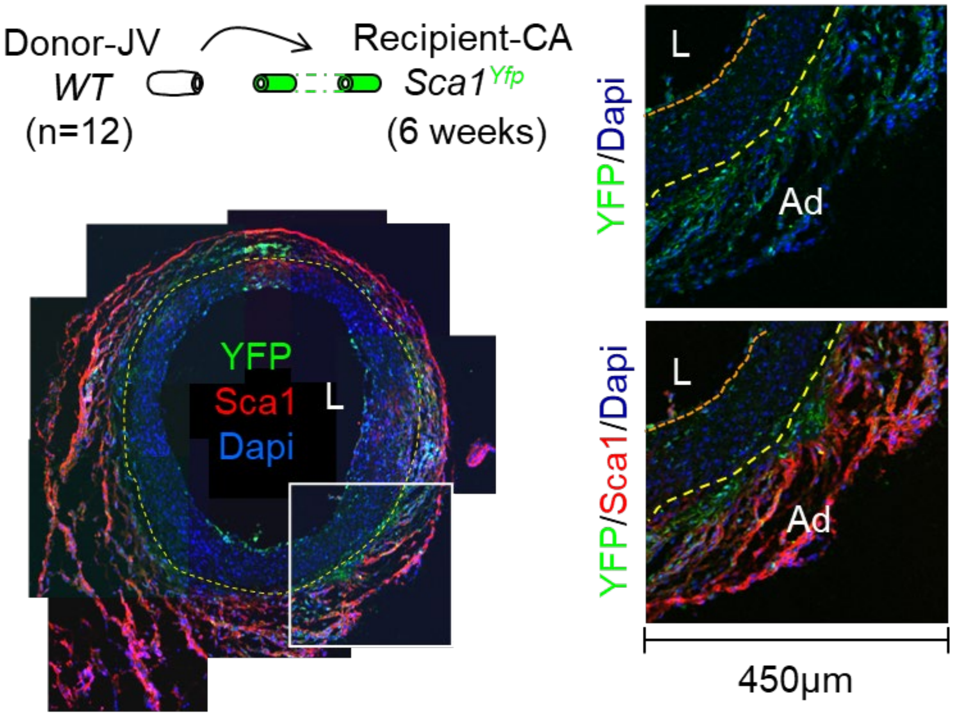


**Figure S7. The Contribution of Recipient Sca1$^+$ Cells to Neointima Formation in VGs.** The representative images of co-staining of YFP with Sca1 in whole cross-sections of wild type (*WT*) jugular vein (JV) grafts transplanted to carotid arteries (CAs) of *Sca1*$^{Yfp}$ mice in which Sca1$^+$ cells are genetically labeled with YFP (n=12). YFP is green; Sca1 is red. L, the lumen; IN, the intima; NI, the neointima; Ad, the adventitia. Dotted orange lines separate the IN and NI, and yellow lines separate the NI and Ad layers.

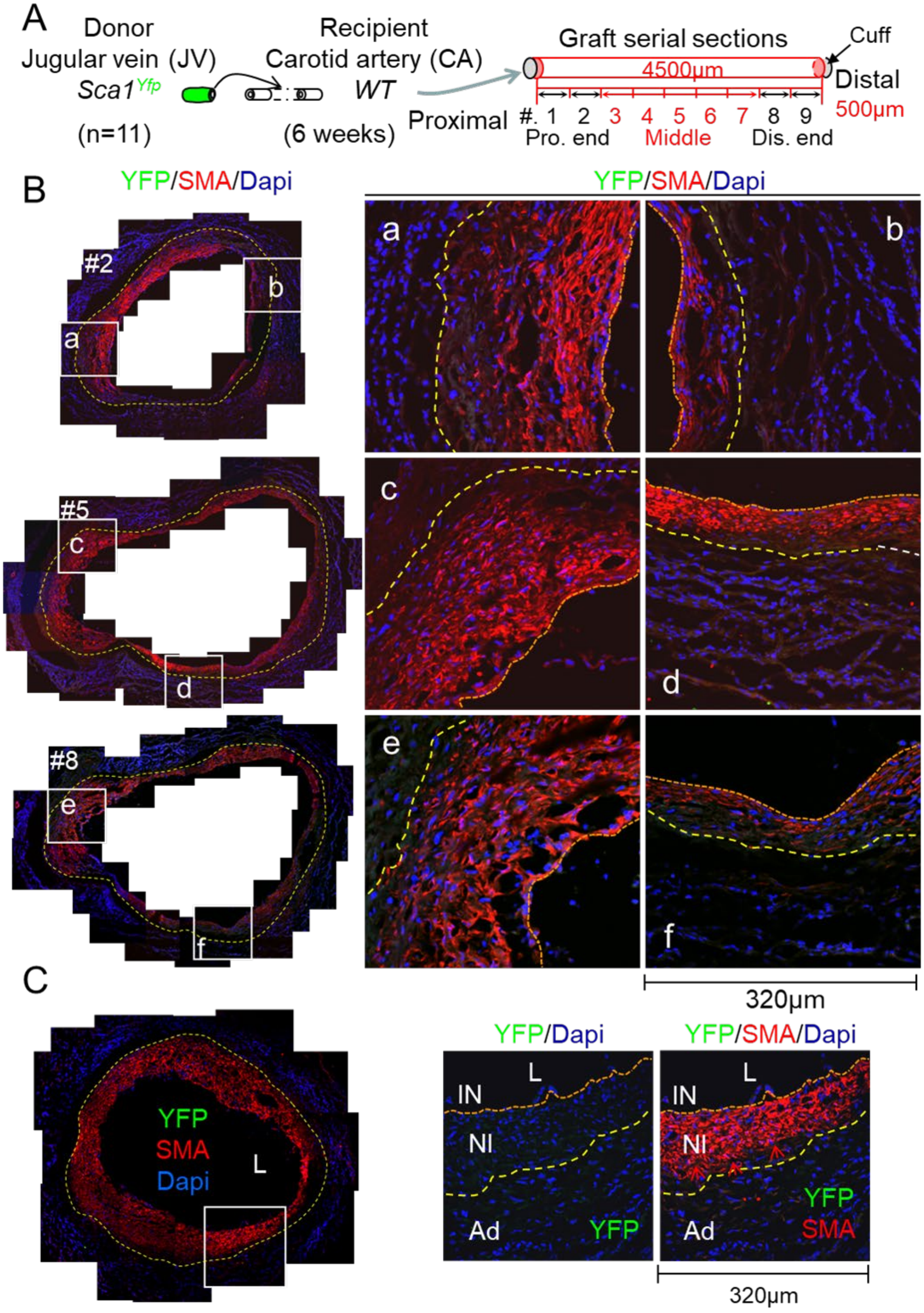


**Figure S8. The Contribution of Donor Venous Sca1⁺ Cells to Neointima Formation in VGs.** (**A**) A Scheme of VG transplantation and cross-tissue sectioning. **Pro.end**, proximal anastomotic end (#1-2); **Middle**, middle body (#3-7); **Dis.end**, distal anastomotic end (#8-9). (**B**)The representative images of co-staining of YFP with SMA in whole cross-sections of in whole cross-sections of *Sca1$^{Yfp}$* reporter JVs grafted into the CAs of *WT* mice (n=11). (**C**) The representative images of co-staining of YFP with SMA in whole cross-sections of *Sca1$^{Yfp}$* reporter JVs grafted into the CAs of *WT* mice for **Figure 1e**.YFP is green; SMA is red. L, the lumen; IN, the intima; NI, the neointima; Ad, the adventitia. Dotted orange lines separate the IN and NI, and yellow lines separate the NI and Ad layers.

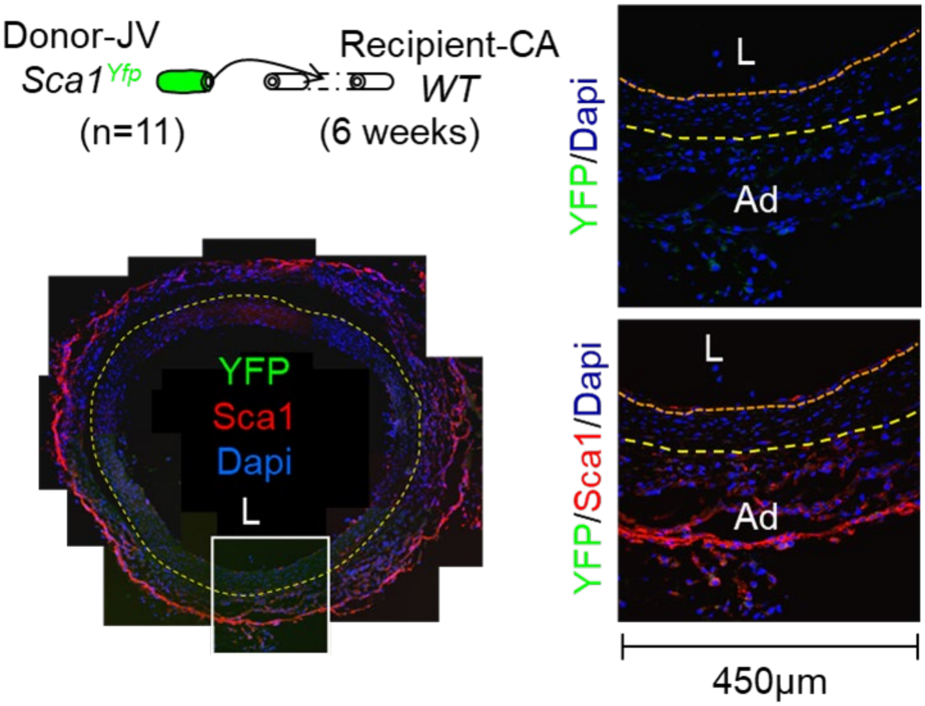


**Figure S9. The Contribution of Donor Venous Sca1$^+$ Cells to Neointima Formation in VGs.** The representative images of co-staining of YFP with Sca1 in whole cross-sections of *Sca1$^{Yfp}$*reporter JVs grafted into the CAs of WT mice for 6 weeks (n=11). YFP is green; Sca1 is red. L, the lumen; IN, the intima; NI, the neointima; Ad, the adventitia. Dotted orange lines separate the IN and NI, and yellow lines separate the NI and Ad layers.

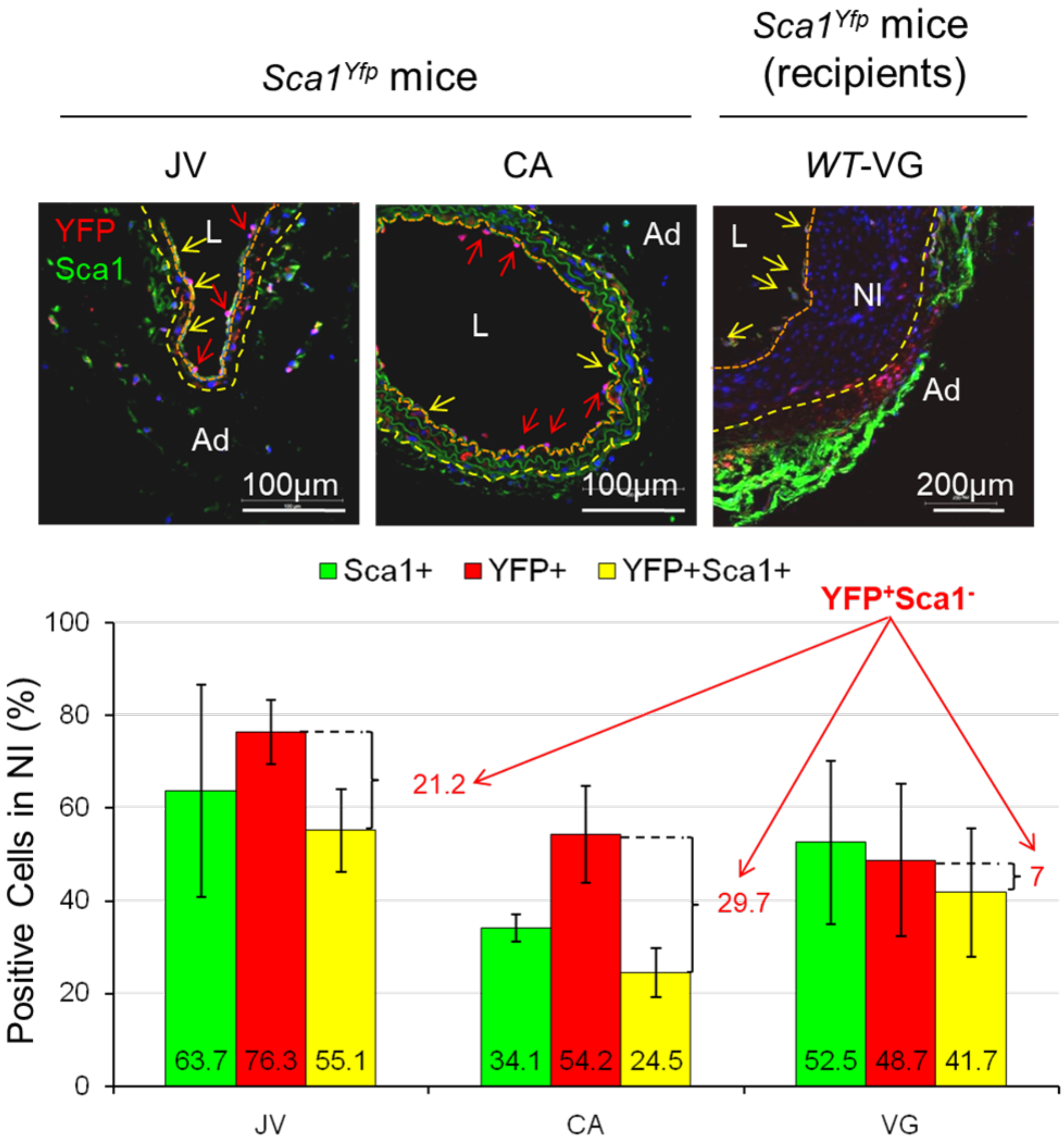


**Figure S10. The endothelialization in normal veins, arteries, and vein grafts.** Jugular veins (JVs) of male *WT* mice at age of 3 months were transplanted into carotid arteries (CAs) of*Sca1-Cre::Rosa26$^{floxedStop}$eYfp* (*Sca1$^{Yfp}$*) mice for 6 weeks. The vein grafts (VGs), normal JVs and CAs of Sca1Yfp mice were harvested for immunofluorescence staining of YFP and Sca1 (n=3). The representative immunofluorescence staining of YFP and Sca1 in JVs, CAs and VGs were shown in the top panel and quantified results are shown in lower panel. YFP is red; Sca1 is green; Red arrows indicate YFP$^{+}$Sca1$^{-}$ cells. Yellow arrows indicate YFP$^{+}$Sca1$^{+}$ cells. L, the lumen; IN, the intima; NI, the neointima; Ad, the adventitia. Dotted orange lines separate the IN and NI, and yellow lines separate the NI and Ad layers. Data were showed as means ± SD (5 whole vessel cross-sections per mice, 15 whole vessel cross-sections were analyzed).

**3. Genetic Ablation of Sca1⁺ Cells in vivo**

*Sca1-Cre::Rosa26$^{floxedStop}$eYfp::Rosa26$^{floxedStop}$DTR* (Cre-inducible transgenic expression of a diphtheria toxin receptor)[10] mice were generated by crossing *Sca1-Cre::Rosa26$^{floxedStop}$eYfp* and *Rosa26$^{floxedStop}$DTR* mice, which render Sca1$^{+}$ cells sensitive to an optimized regiment of diphtheria toxin (DT) treatment, thereby selectively depleting vascular Sca1$^{+}$ cells in mice (**Figures S11-12, and Table S6**).The recipient Sca1$^{+}$ cells were accumulated in the adventitia but not in the intima of VGs at 21 days after transplantation (**Figure S12B**).

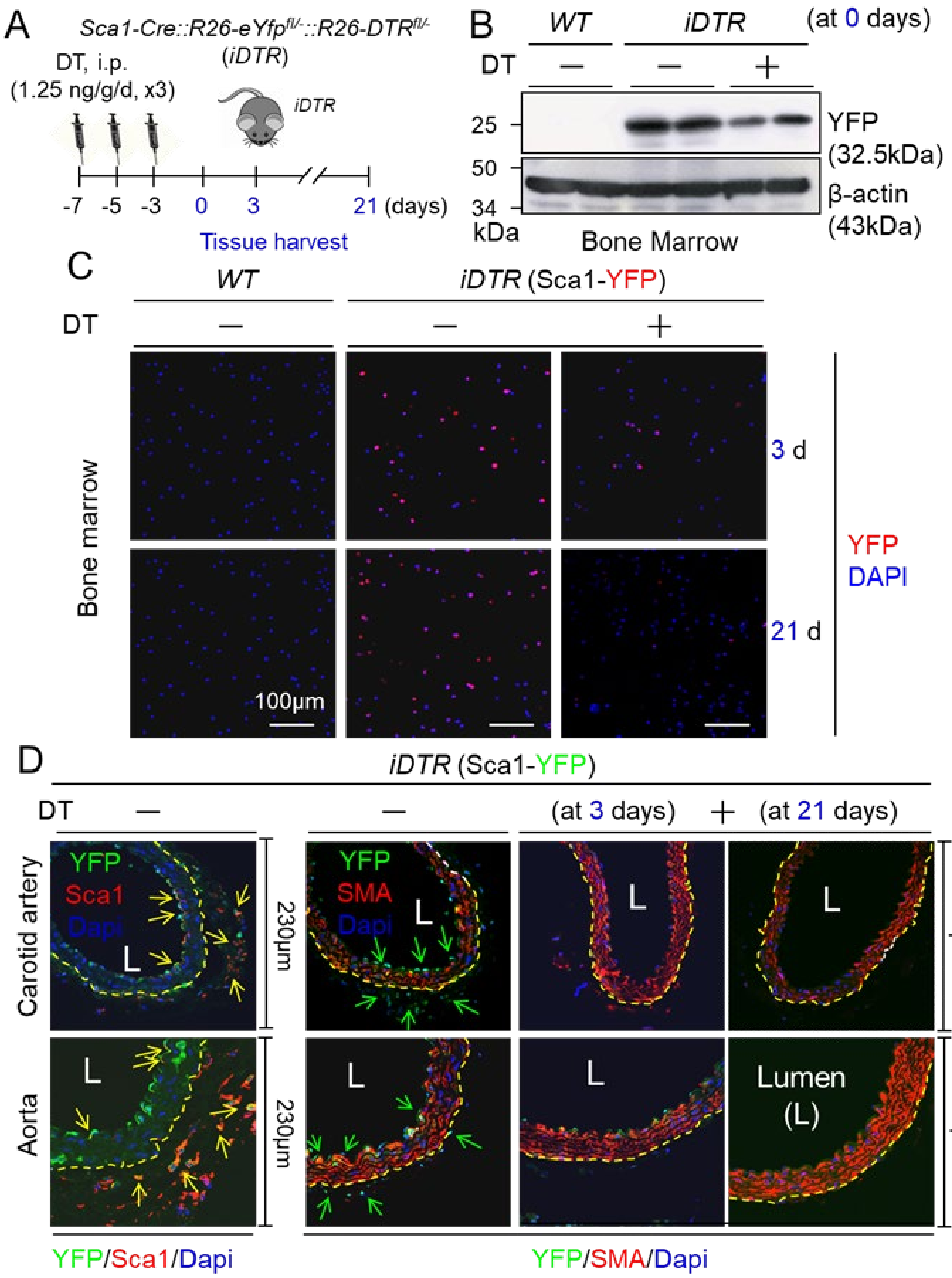

**Figure S11. Establishing a Regiment of Diphtheria Toxin (DT) Treatment for Ablating Vascular Sca1$^+$ Cells in Adult *Sca1-Cre::Rosa26$^{floxedStop}$eYfp::Rosa26$^{floxedStop}$DTR* (*iDTR*) Mice.** Male 12-week-old *iDTR* mice were i.p. injected with DT once every two days at a dose of 1.25 ng/kg/d for 3 times, followed 3-day washout period, and then subject to tissue harvest for Western blot analysis and immunochemical staining. (**A**) A scheme of experimental procedures. (**B**) Western blot analysis of YFP expression in bone marrow of *iDTR* mice at 0 day after receiving the optimized regiment of DT treatment (n=2). (**C**) Immunofluorescence staining of YFP positive cells (Sca1$^+$ cells) in bone marrow of *Idtr* mice at 3 and 21 days after receiving the optimized regiment of DT treatment (n=3). (**D**) The Impact of genetic ablation of vascular Sca1$^+$cells on arterial structure at basal condition. Representative immunofluorescence staining of YFP, Sca1, and SMA in arteries of the *iDTR* mice after receiving the optimized regiment of DT treatment at basal condition (n=3, 3 whole cross-sections per mice).

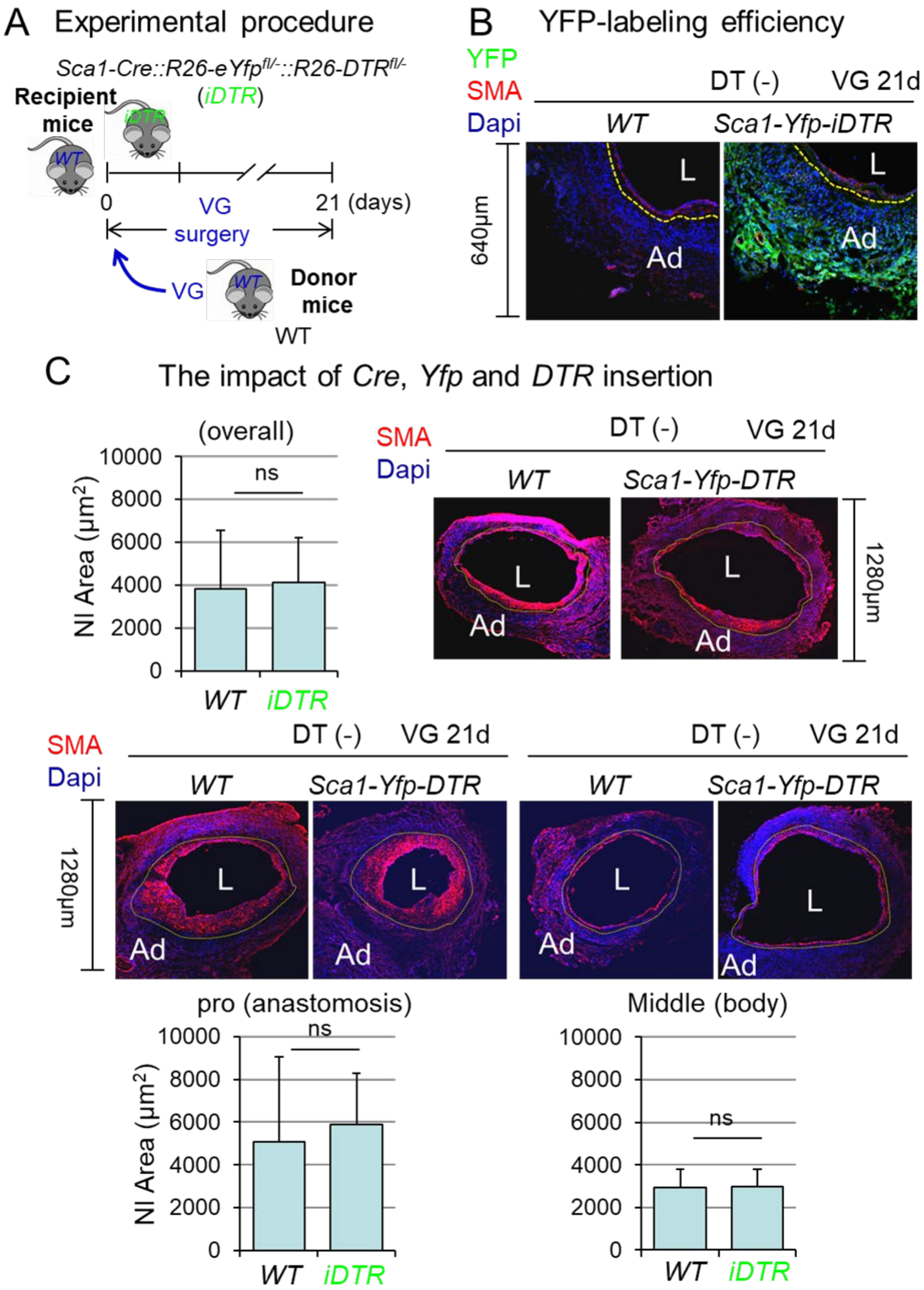


**Figure S12. The Impact of *Cre*, *Yfp*, and *DTR* Gene Insertions in Recipient *Sca1-Cre::Rosa26$^{floxedStop}$eYfp::Rosa26$^{floxedStop}$DTR* mice on Vascular Remodeling of Vein Grafts of Wild Type (*WT*) Donor.** Jugular veins of male 3-month old *WT* mice were transplanted into carotid arteries of *Sca1-Cre::Rosa26$^{floxedStop}$eYfp::*

*Rosa26$^{floxedStop}$DTR*(*Sca1-Yfp-DTR* or *iDTR*) without diphtheria toxin (DT) treatment for 21 days (n=4~5).(**A**) A scheme of experimental procedures. VG, vein graft. (**B**) YFP labeling efficiency. Representative immunofluorescence staining of YFP and SMA in *WT* jugular veins transplanted into the *iDTR* mice without DT treatment at 21 days after transplantation (n=4~5). YFP$^+$ cells (recipient Sca1$^+$ cells) are mainly in the adventitia but hardly observable in the intima of VGs. L. the lumen; Ad, the adventitia. (**C**) Remodeling of *WT* VGs transplanted into carotid arteries of *iDTR* mice without DT treatment (n=4~5). There are negligible effects of *Cre*, *Yfp*, or *DTR* insertions on neointima formation in VGs. Three whole cross sections from each VG were analyzed. L, the lumen; Ad, the adventitia. Dotted yellow lines separate the NI and Ad layers. Data are means ± SD. ns, non-significant. The statistical analysis is carried out by Student t test.

**4. Delivery Route-Dependent Therapeutic Efficacy of Simvastatin in Suppressing VG Intimal Hyperplasia**

The effects of Simvastatin via different delivery routes on vein graft (VG) intimal hyperplasia were compared in mice. The therapeutic efficacy of peri-vascular delivery of simvastatin for 3 days after transplantation is as effective as the statin administration 3 days prior to transplantation;[11] however, simvastatin administration started 3 days after transplantation fails to inhibit the neointima formation (**Figure S13**).

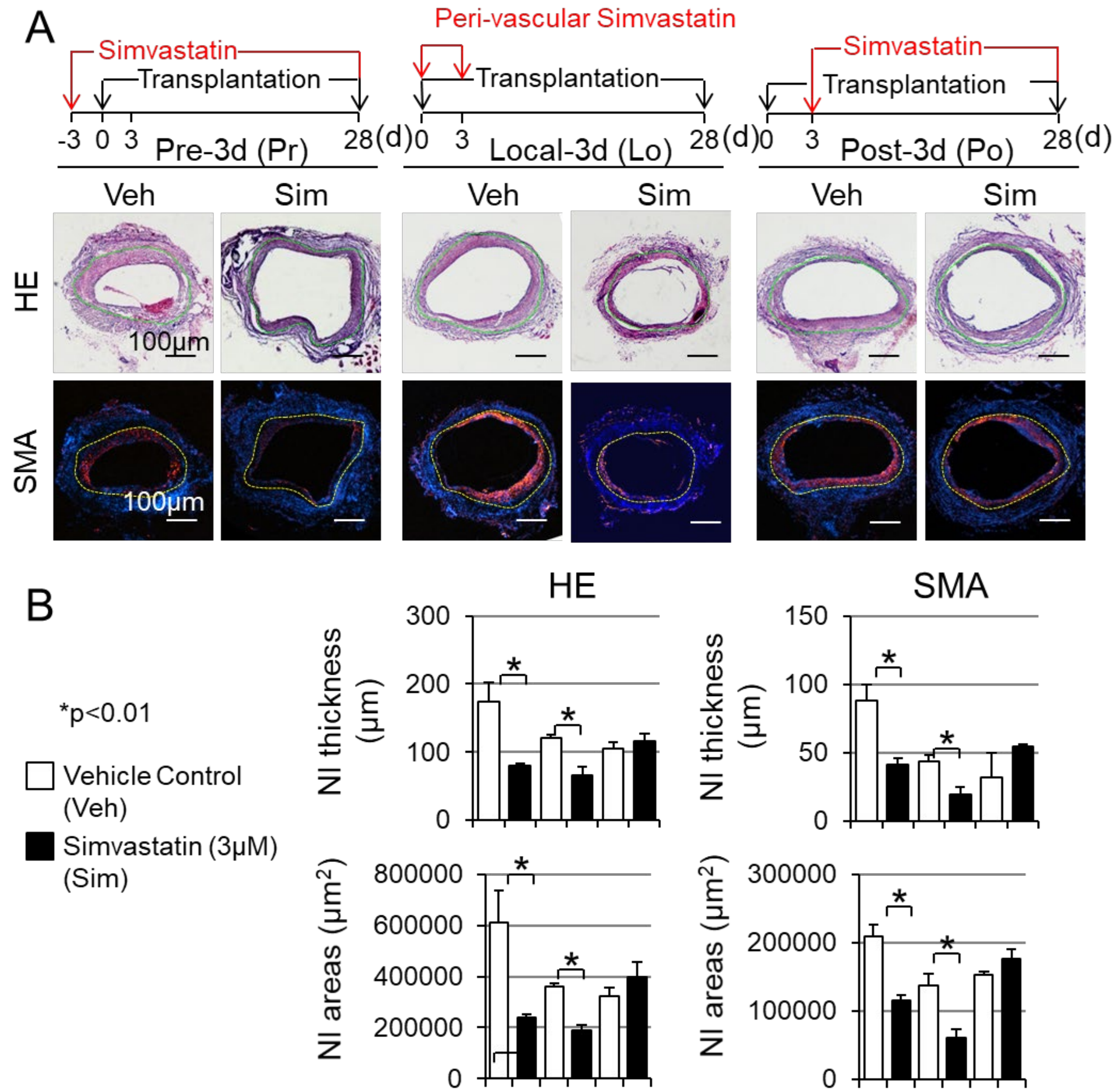


**Figure S13. Delivery Route-Dependent Effects of Simvastatin on Vein Graft (VG) Remodeling.** The therapeutic efficacy of simvastatin via different delivery routes were determined in mouse model of vein graft intimal hyperplasia (*WT* to *WT* transplantation). (**A**) Schematic presentations of simvastatin deliveries and representative HE and immunofluorescence staining of whole VG cross-sections at 28 days after transplantation (n=4). Three routes of simvastatin administration:(1) intragastric administration of simvastatin (1.6 mg/kg/d) daily from 3 days before transplantation until the endpoint; (2) intragastric administration of simvastatin (1.6 mg/kg/d) daily starting 3 days after transplantation until the endpoint; 3) peri-vascular delivery of simvastatin with pluronic-127 gel (3 μM in 50 μL) immediately after transplantation, which could locally release the drug up to 3 days. (**B**) Quantified neointima (NI) areas, NI thickness, and SMA positive areas and thickness. Nine cross-sections of each VG were subject to the analysis (n=4). **P*<0.05 vs. vehicle control (Veh). Dotted yellow lines separate the NI and Ad layers. Data are means ± SD. ns, non-significant. The statistical analysis is carried out by Student t test.

**5. Selectively Effects of Simvastatin on Adventitial Sca1⁺ Vascular Stem Cell (Sca1⁺AdVSC) Growth and Sca1⁺AdVSC-Mediated Medial SMC Migration and Proliferation**

Simvastatin at doses of 0.25 – 0.5 μM which inhibited proliferation of mouse aortic $Sca1^+$ VSCs (**Figure 1h**) did not affect proliferation of mouse aortic SMCs in vitro (**Figure S14**). $Sca1^+$ cells isolated from mouse aortic adventitia at passage 1 (P1) did not express embryonic stem cell markers OCT4 but mesenchymal stem cell markers CD44 and CD45 and vascular stem cell markers NFM and SOX10 as well as SMC markers SMA, Smoothelin, SM22, and CNN1, and EC markers TIE2, CD31, and vWF (**Figure S15A**). These cells up to passage 10 can be differentiated into chondrocytes, adipocytes, and osteoblasts (**Figure S15A**). These results reveal that the adventitial $Sca1^+$ cell are vascular stem cells (VSCs) ($Sca1^+$AdVSCs), which are primarily functioning as vascular progenitor cells. In addition, the conditioned media from sub-cultured $Sca1^+$AdVSCs increased aortic SMC migration and proliferation (**Figure S15B-D**), indicating that activated adventitial $Sca1^+$ VSCs can promote the media SMC dedifferentiation towards the synthetic type of SMCs via paracrine regulation. Notably, simvastatin decreased the effect of $Sca1^+$AdVSC conditioned media on SMC proliferation and migration (**Figures S15C, S15D**). On the other hand, simvastatin promoted $Sca1^+$AdVSC differentiation to ECs and SMCs while inhibiting $Sca1^+$AdVSC differentiation to chondrocytes, adipocytes, and osteoblasts in vitro (**Figure S16**), revealing a simvastatin-induced naïve state of $Sca1^+$AdVSCs.

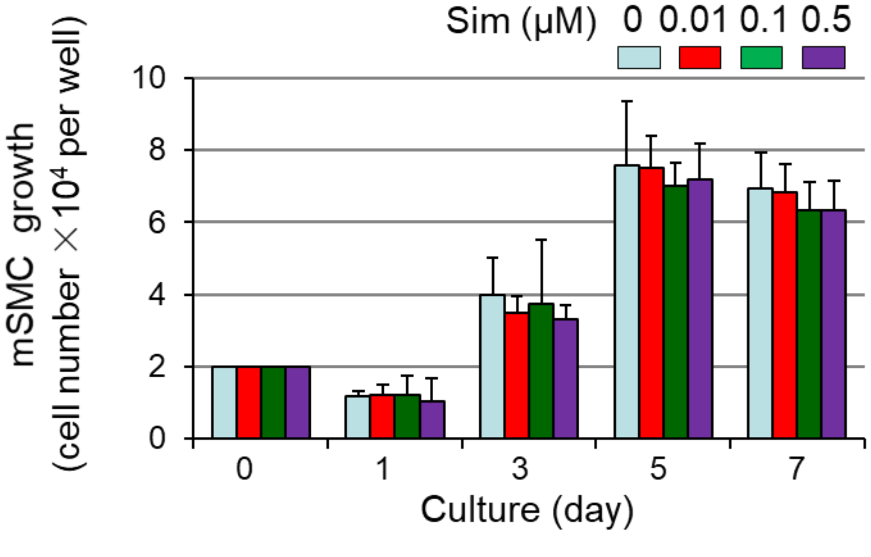


**Figure S14. Effect of Simvastatin on Proliferation of cultured mouse aortic SMCs.** Growth curve of mouse aortic SMCs cultured in DMEM + 10%FBS with or without simvastatin (Sim, 0, 0.01, 0.1, 0.5μM) for 7 days (n=4). Simvastatin at doses of 0.25 – 0.5 μM did not inhibit proliferation of mouse aortic SMCs in vitro.

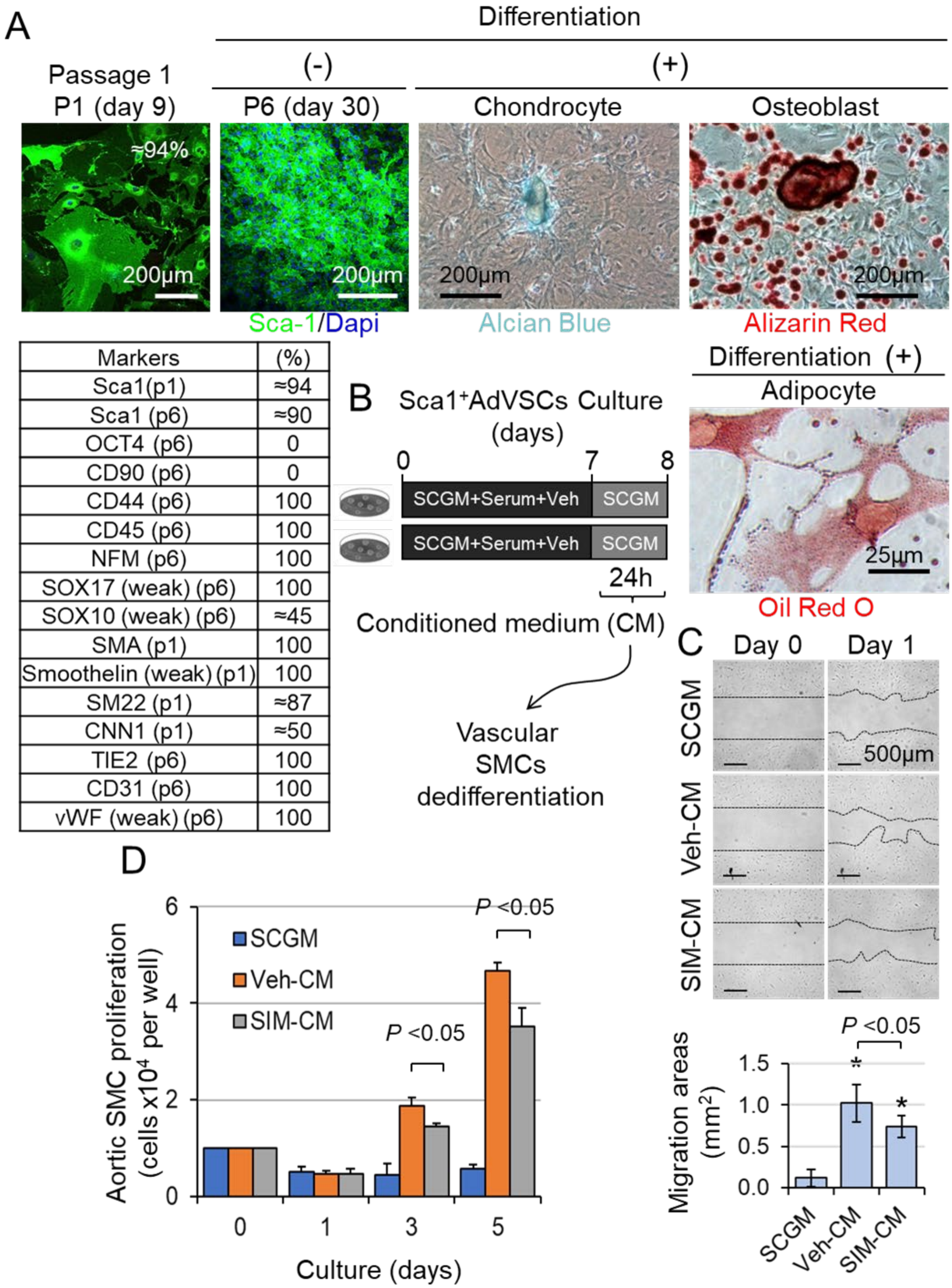


| Markers | (%) |
|---|---|
| Sca1(p1) | ≈94 |
| Sca1 (p6) | ≈90 |
| OCT4 (p6) | 0 |
| CD90 (p6) | 0 |
| CD44 (p6) | 100 |
| CD45 (p6) | 100 |
| NFM (p6) | 100 |
| SOX17 (weak) (p6) | 100 |
| SOX10 (weak) (p6) | ≈45 |
| SMA (p1) | 100 |
| Smoothelin (weak) (p1) | 100 |
| SM22 (p1) | ≈87 |
| CNN1 (p1) | ≈50 |
| TIE2 (p6) | 100 |
| CD31 (p6) | 100 |
| vWF (weak) (p6) | 100 |

**Figure S15. Inhibitory Effects of Simvastatin on Adventitial Sca1+ Vascular Stem Cell (Sca1+AdVSC) Activation and Paracrine Enforcement of Medial SMC**

**Dedifferentiation.** (**A**) **Characterization of Sca1$^+$AdVSC stemness**. The expression of stem cell markers in cultured Sca1$^+$AdVSCs isolated from mouse aortic adventitial layers at different passages was determined. The representative images of immunofluorescence staining for Sca1+ AdVSC's differentiation to chondrocytes (Alcian blue), adipocytes (Oil Red O), and osteoblasts (Alizarin Red) are from 3 separated experiments. (**B**) A scheme of Sca1$^+$AdVSC conditioned medium-induced medial SMC dedifferentiation. (**C**) The effect of simvastatin on Sca1$^+$AdVSC conditioned medium-induced migration of medial SMCs. Sub-cultured mouse aortic SMCs at a confluent state were subject to scratch assay. Plain stem cell growth medium with 0.1% FBS is used as the control. The effects of conditioned media (0.1% FBS) of Sca1$^+$AdVSCstreated with vehicle (Veh) and simvastatin (0.5 µM Sim) on aortic SMC migration were determined as indicated (n=4). *$P$< 0.05 vs SCGM group; $P$< 0.05 between indicated groups. (**D**)The effect of simvastatin on Sca1$^+$AdVSC conditioned medium-induced proliferation of medial SMCs. Mouse aortic SMCs were cultured in SCGM and conditioned media of Sca1$^+$AdVSCs treated with Veh or Sim (0.5 µM) supplemented with 1% FBS for 5 days (n=4). $P$< 0.05 between indicated groups.

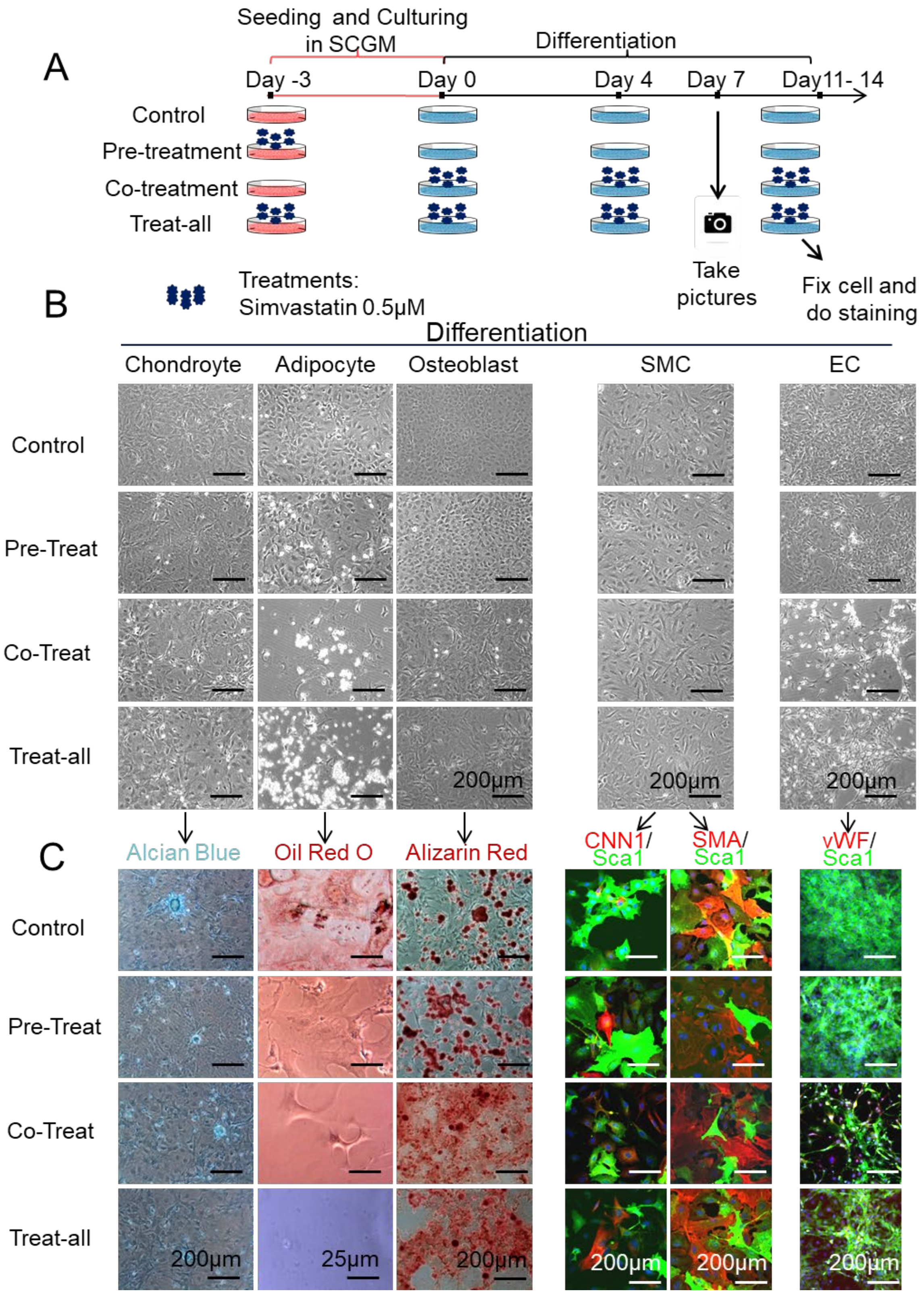


**Figure S16. The Effect of Simvastatin on Adventitial Sca1$^{+}$ Vascular Stem Cell (Sca1$^{+}$AdVSC) Differentiation.** (**A**) Schematic diagram of differentiation for mouse

Sca1$^+$AdVSCs with or without simvastatin (0.5µM) treatment. Simvastatin was administrated for 3 days alone prior to the induced differentiation, i.e., pretreatment; added into differentiation media until end of the differentiation, i.e., cotreatment; treated 3 days before the differentiation until end of the differentiation, i.e., all-treat. (**B**) The representative images of bright field staining for Sca1$^+$AdVSC's differentiation to chondrocytes (Alcian blue), adipocytes (Oil Red O), and osteoblasts (Alizarin Red) in bright field. (**C**) The representative images of immunofluorescence staining for Sca1$^+$AdVSC's differentiation to SMC (CNN1/SMA-red and Sca1-green) and EC (vWF-red and Sca1-green).

**Table S1. Cellular Dynamics of VSCs, SMCs, and ECs in Vein Graft Remodeling.** Vein graft (VG) transplantation was carried out using male 3-month-old WT mice in a C57BL/6J genetic background. VGs were harvested at 0, 1, 3, 7, 14, 42 days after transplantation and subject to morphological analyses as described in Methods (n=8). The percentages of cells positive for Sca1, Gli1, CD34, Tunel, and Ki67 in normal jugular veins and VGs were quantified (IN, intima; M, media; NI, neointima; Ad, adventitia) (n=8). Three cross tissue sections from each vessel were analyzed for each biomarker. Data are means ± SD. d, day. **NOTE**: 0d-VGs, IN+NI = IN+M (no NI formation); transplanted-VGs, IN+NI = IN+NI+M (It is hard to differentiate NI from M in remodeling VGs in mice, NI here includes both NI and M.).

| **Sca1 (%)** | **IN+NI** | **Ad** |
|---|---|---|
| 0d | 39.47±18.11 | 37.39±19.32 |
| 1d | 0±0 | 19.1±2.11 |
| 3d | 0±0 | 22.73±6.43 |
| 7d | 11.16±7.53 | 28.65±4.93 |
| 14d | 11.07±3.03 | 35.05±11.57 |
| 42d | 5.83±2.65 | 74.19±4.5 |
| **Tunel (%)** | **IN+NI** | **Ad** |
| 0d | 0±0 | 0±0 |
| 1d | 0.8±0.09 | 0.36±0.05 |
| 3d | 0.89±0.02 | 0.52± |
| 7d | 0.03±0.01 | 0.01±0.03 |
| 14d | 0.01±0 | 0±0 |
| 42d | 0±0 | 0±0 |
| **Ki67 (%)** | **IN+NI** | **Ad** |
| 0d | 0±0 | 0±0 |
| 1d | 0±0 | 0.08±0.04 |
| 3d | 0±0 | 0.13±0.02 |
| 7d | 0.04±0 | 0.54±0.03 |
| 14d | 0.54±0.05 | 0.13±0.06 |
| 42d | 0.01±0.01 | 0.02±0.01 |
| **CD34 (%)** | **IN+NI** | **Ad** |
| 0d | 43.56±9.11 | 81.13±31.42 |
| 1d | 0±0 | 14.73±7.38 |
| 3d | 0±0 | 16.47±5.9 |
| 7d | 1.05±2.35 | 22.98±13.09 |
| 14d | 1.36±2.1 | 29.52±11.21 |
| 42d | 6.62±2.8 | 48.35±15.1 |
| **Gli1 (%)** | **IN+NI** | **Ad** |
| 0d | 0±0 | 76.44±12.84 |
| 1d | 0±0 | 0±0 |
| 3d | 0±0 | 0±0 |
| 7d | 0±0 | 0±0 |
| 14d | 0±0 | 0±0 |
| 42d | 0±0 | 77.74±15.65 |

**Table S2. The Heterogeneity of Vascular Stem Cells in Vein Graft Remodeling.** Vein graft (VG) transplantation was carried out using male 3-month-old WT mice in a C57BL/6J genetic background. VGs were harvested at 0, 1, 3, 7, 14, 42 days after transplantation and subject to morphological analyses as described in Methods (n=8). The percentages of cells double positive for Sca1 and Gli1 ($Sca1^+Gli1^+$), $Sca1^+CD34^+$, $Sca1^+SMA^+$, or $Sca1^+vWF^+$ in normal jugular veins and VGs were quantified (IN, intima; M, media; NI, neointima; Ad, adventitia) (n=8). Three cross tissue sections from each vessel were analyzed for each biomarker. Data are means ± SD. d, day. 0d-VGs, IN+NI = IN+M (no NI formation); transplanted-VGs, IN+NI = IN+NI+M (It is hard to differentiate NI from M in remodeling VGs in mice, NI here includes both NI and M.).

| **$Sca1^+Gli1^+$ (%)** | **IN+NI** | **Ad** |
|---|---|---|
| 0d | 0±0 | 33.86±15.06 |
| 1d | 0±0 | 0±0 |
| 3d | 0±0 | 0±0 |
| 7d | 0±0 | 0±0 |
| 14d | 0±0 | 0±0 |
| 42d | 0±0 | 65.18±7.83 |
| **$Sca1^+CD34^+$ (%)** | **IN+NI** | **Ad** |
| 0d | 31.23±12.35 | 45.16±19.97 |
| 1d | 0±0 | 1.68±1.2 |
| 3d | 0±0 | 6.14±3.66 |
| 7d | 1.05±2.35 | 7.79±7.94 |
| 14d | 0±0 | 19.47±7.05 |
| 42d | 1.49±2.14 | 40.31±15.75 |
| **$Sca1^+SMA^+$ (%)** | **IN+NI** | **Ad** |
| 0d | 0±0 | 0±0 |
| 1d | 0±0 | 0±0 |
| 3d | 0±0 | 0±0 |
| 7d | 4.34±1.78 | 6.63±1.42 |
| 14d | 9.07±2.04 | 12.69±3.82 |
| 42d | 6.72±0.28 | 5.52±3.31 |
| **$Sca1^+vWF^+$ (%)** | **IN+NI** | **Ad** |
| 0d | 18.98±5.91 | 0±0 |
| 1d | 0±0 | 0±0 |
| 3d | 0±0 | 0±0 |
| 7d | 0±0 | 0±0 |
| 14d | 0±0 | 0±0 |
| 42d | 4.67±4.2 | 0±0 |

**Table S3. Tracing the Fate of Recipient Sca1⁺ Cells in VG Remodeling.** Jugular veins of male wild type (*WT*) mice were transplanted into carotid arteries (CA) of male littermates of *Sca1-Cre::Rosa26$^{floxedStop}$eYfp* (*Sca1$^{YFP}$*) mice at age of 3-months for 6 weeks (n=12). **VG**, vein graft; **IN**, the intima; **NI**, the neointima; **Ad**, the adventitia

| | ***WT*** VGto***Sca1$^{Yfp}$*** CA (n=12) | | |
|---|---|---|---|
| | Carotid Artery (CA) of *Sca1$^{Yfp}$* mice at the baseline | | |
| (%) | Intima (IN) | Media (M) | Adventitia (Ad) |
| $SMA^+$/Dapi | 0±0 | 87.30±4.46 | 1.56±1.48 |
| $Sca1^+$/Dapi | 34.09±2.94 | 0±0 | 62.63±5.49 |
| $YFP^+$/Dapi | 54.23±10.36 | 8.92±1.82 | 48.89±7.18 |
| $YFP^+SMA^-$/Dapi | 0±0 | 8.78±1.64 | 0.38±0.86 |
| $YFP^+Sca1^+$/Dapi | 24.5±5.32 | 0±0 | 40.58±6.67 |
| | *WT* VGs at 6 weeks after transplantation | | |
| $SMA^+$/Dapi (%) | Intima (IN) | Neointima (NI) | Adventitia (Ad) |
| Proximal end | 0±0 | 69.72±4.77 | 6.74±5.49 |
| Middle body | 0±0 | 63.70±6.21 | 3.94±4.04 |
| Distal end | 0±0 | 64.27±9.13 | 19.99±2.89 |
| $YFP^+$/Dapi (%) | IN | NI | Ad |
| Proximal end | 35.98±18.67 | 8.51±10.05 | 76.05±7.32 |
| Middle body | 45.49±18.26 | 14.03±10.97 | 80.00±6.22 |
| Distal end | 37.78±11.44 | 16.62±7.50 | 78.18±5.43 |
| $SMA^+YFP^+$/Dapi (%) | IN | NI | Ad |
| Proximal end | 0±0 | 1.65±2.06 | 4.07±2.99 |
| Middle body | 0±0 | 0.49±0.55 | 1.21±1.46 |
| Distal end | 0±0 | 2.62±1.81 | 8.72±1.75 |

**Table S4. The Fate of Donor Venous Sca1⁺ Cells in Vein Graft Remodeling.** The jugular veins (n=12) of male 3-month-old *Sca1-Cre::Rosa26$^{floxedStop}$eYfp* (*Sca1-Yfp*) littermates were transplanted into carotid arteries of male wild type (*WT*) mice age of 3 months for 6 weeks. VG, vein graft; IN, the intima; NI, the neointima; Ad, the adventitia.

| | ***Sca1$^{Yfp}$***VGto***WT***CA | | |
|---|---|---|---|
| | Jugular vein (JV) graft at the baseline | | |
| | Intima (IN) | Media (M) | Adventitia (Ad) |
| $SMA^+$/Dapi(%) | 0±0 | 95.71±10.29 | 1.25±1.82 |
| $Sca1^+$/Dapi(%) | 61.71±5.33 | 0±0 | 74.3±6.35 |
| $YFP^+$/Dapi(%) | 76.28±7 | 2.25±2.18 | 50.6±0.89 |
| $YFP^+SMA^-$/Dapi(%) | 0±0 | 2.25±2.18 | 0.45±1.01 |
| $YFP^+Sca1^+$/Dapi(%) | 55.13±8.96 | 0±0 | 46.62±1.38 |
| | ***Sca1-Yfp*** VG at 6 weeks after transplantation | | |
| $SMA^+$/Dapi (%) | Intima (IN) | Neointima (NI) | Adventitia (Ad) |
| Proximal end | 0±0 | 79.09±19.73 | 0.99±0.83 |
| Middle body | 0±0 | 76.14±8.04 | 0.99±1.60 |
| Distal end | 0±0 | 75.42±10.81 | 1.17±2.89 |
| $YFP^+$/Dapi (%) | I | NI | Ad |
| Proximal end | 0±0 | 0±0 | 0±0 |
| Middle body | 0±0 | 0±0 | 0±0 |
| Distal end | 0±0 | 0±0 | 0±0 |
| $SMA^+YFP^+$/Dapi (%) | I | NI | Ad |
| Proximal end | 0±0 | 0±0 | 0±0 |
| Middle body | 0±0 | 0±0 | 0±0 |
| Distal end | 0±0 | 0±0 | 0±0 |

**Table S5. Primers for Genotyping.**

| Mouse strain | Primers | Reaction conditions | Productions |
|---|---|---|---|
| *Sca-1-Cre* mice | Primer 1 (PB003):<br>5'TGCAACGAGTGATGAGGTTCGC3'<br><br>Primer 2 (PB004):<br>5'GATCCTGGCAATTTCGGCTATACG3'<br><br>Primer 3 (1136):<br>5'GGTTTCTATTGCTACCAAGAGACAT3'<br><br>Primer 4 (1044):<br>5'TGCACCAAACCCTGGACTAAGCAT3' | Cycling:<br>95°C, 30 sec;<br>95°C, 30 sec;<br>58°C, 30 sec;<br>72°C, 1 min;<br>Repeat 35 cyc;<br>72°C, 6 min;<br>4°C, hold. | Internal positive control:<br>407 bp<br><br>Cre:<br>490 bp |
| *Rosa26* *Floxed-StopeYFP* mice Stock Number: 006148 (JAX) | Primer1 (oIMR8546):<br>5'GGAGCGGGAGAAATGGATATG3'<br><br>Primer2 (oIMR8545):<br>5'AAAGTCGCTCTGAGTTGTTAT3'<br><br>Primer3 (oIMR4982):<br>5'AAGACCGCGAAGAGTTTGTC3' | Cycling:<br>95°C, 5 min;<br>95°C, 30 sec;<br>58°C, 30 sec;<br>72°C, 45 sec;<br>Repeat 35 cyc;<br>72°C, 5 min;<br>4°C. | Mutant:<br>320 bp<br><br>Heterozygote:320 bp and 603 bp<br><br>Wild type:<br>603 bp |
| *Rosa26* *Floxed-StopDTR* mice Stock Number: 007900 (JAX) | Primer1 (oIMR8546):<br>5'GGAGCGGGAGAAATGGATATG3'<br><br>Primer2 (oIMR8545):<br>5'AAAGTCGCTCTGAGTTGTTAT3'<br><br>Primer3 (oIMR8052):<br>5'GCGAAGAGTTTGTCCTCAACC3' | Cycling:<br>95°C, 2 min;<br>95°C, 20 sec;<br>65°C,<br>0.5°C per cycle decrease;<br>68°C, 10 sec;<br>Repeat 2-4 10 cyc;<br>95°C, 15 sec;<br>60°C, 15 sec;<br>72°C, 10 sec;<br>Repeat 6-8 28cyc;<br>72°C, 5 min;<br>10°C, hold | Mutant:<br>300 bp<br><br>Heterozygote: 300 bp and 603 bp;<br><br>Wild type:<br>603 bp |

**Table S6. Titration of Diphtheria Toxin (DT) Doses for Ablating Sca1$^{+}$ Cells in Adult *Sca1-Cre::Rosa26$^{floxedStop}$eYfp:: Rosa26$^{floxedStop}$DTR* (*iDTR*) Mice.** The toxicity of different doses of DT was tested in *iDTR* mice. Male or female 12-week-old *iDTR* mice were intraperitoneally (i.p.) injected with DT once every two daysat doses as indicated. **Exp.**, experiment groups. Male (♂); female (♀).

| Exp. | DT | *iDTR* mice | Injection (i.p.) | Outcome |
|---|---|---|---|---|
| #1. | 50 (ng/g/d) | 3-month-old ♀ mice (n=2) | One injection, i.e., x1 | All died within 12 h post-injection |
| #2. | 10 (ng/g/d) | 3-month-old ♀ mice (n=2) | One injection, i.e., x1 | All died at 2$^{nd}$ day post-injection |
| #3. | 5 (ng/g/d) | 3-month-old ♀ (n=1) and ♂ (n=1) mice | Once every 2 days, 2 times, i.e., x2 | All died at 3$^{rd}$ day post-injection |
| #4. | 2.5 (ng/g/d) | 3-month-old ♀ (n=1) and ♂ (n=1) mice | Once every 2 days, 2 times, i.e., x2 | The female died at 1$^{st}$ day and the male died at 3$^{rd}$ day post-injection |
| #5. | 1.25 (ng/g/d) | 3-month-old ♂ mice (n=2) | Once every 2 days, 3 times, i.e., x3 | All survived |
